# A Multi-Phenotype System to Discover Therapies for Age-Related Dysregulation of the Immune Response to Viral Infections

**DOI:** 10.1101/2020.07.30.223875

**Authors:** Brandon White, Ben Komalo, Lauren Nicolaisen, Matt Donne, Charlie Marsh, Rachel M. DeVay, An M. Nguyen, Wendy Cousin, Jarred Heinrich, William J. Van Trump, Tempest Plott, Colin J. Fuller, Dat Nguyen, Daniel Chen, Delia Bucher, Sabine Tyrra, Laura Haynes, George Kuchel, Jorg Goronzy, Anis Larbi, Tamas Fulop, Diane Heiser, Ralf Schwandner, Christian Elabd, Ben Kamens

## Abstract

Age-related immune dysregulation contributes to increased susceptibility to infection and disease in older adults. We combined high-throughput laboratory automation with machine learning to build a multi-phenotype aging profile that models the dysfunctional immune response to viral infection in older adults. From a single well, our multi-phenotype aging profile can capture changes in cell composition, physical cell-to-cell interaction, organelle structure, cytokines, and other hidden complexities contributing to age-related dysfunction. This system allows for rapid identification of new potential compounds to rejuvenate older adults’ immune response. We used our technology to screen thousands of compounds for their ability to make old immune cells respond to viral infection like young immune cells. We observed beneficial effects of multiple compounds, of which two of the most promising were disulfiram and triptonide. Our findings indicate that disulfiram could be considered as a treatment for severe coronavirus disease 2019 and other inflammatory infections.

## INTRODUCTION

Aging increases risk of infection progressing to severe disease and death.^1^ The cause of this increased risk is not well understood, but it is known that the immune system undergoes many changes as it ages. Some changes impact the innate immune system, such as the dysregulation of inflammatory responses—otherwise known as “inflamm-aging”—which may disrupt innate antiviral responses by increasing the resting level of circulating proinflammatory cytokines.^2–4^ Other changes associated with age may impair the adaptive immune system, ranging from relative reductions in naïve T cells and increases in memory/effector T-cell populations to alterations in signaling pathways, epigenetics, and effector functions.^5–11^ Collectively, these age-related alterations in the immune system are referred to as “immunosenescence.”^1,6^

Yet, while immunosenescence and inflamm-aging have been linked with pathology, both may also have beneficial effects.^2,12^ Maintaining a balance between the beneficial effects of these aging processes while targeting those that are detrimental requires an ability to measure and model a complex interplay of immune states and phenotypes. Such a model could be a powerful system for finding potential therapies for the dysregulated viral response in older adults.

Coronavirus disease 2019 (COVID-19) caused by severe acute respiratory syndrome coronavirus 2 (SARS-CoV-2) exemplifies this increased risk of infection leading to hospitalization or death in older adults.^13–15^ The disease has disproportionally affected older adults who suffer from the highest morbidity and mortality.^13–15^ A study of patients hospitalized with SARS-CoV-2 infection demonstrated that age at admission was associated with a greater mortality risk than other factors such as obesity and chronic cardiac disease (age 70–79 years [HR, 9.6]; age ≥80 years [HR, 13.6]; obesity [HR, 1.4]; chronic cardiac disease [HR, 1.3]).^15^ In China, the case fatality rate was 0.4% for patients age 40–49 years and 15% for patients ≥80 years of age^14^; in Italy, 96% of COVID-19 deaths occurred in people ≥60 years of age, while only 1% of patients under the age of 50 years died.^13^ Some believe that the increased severity of COVID-19 in older adults may be exacerbated by the aging immune system.^16^ The importance of treating the host immune response was recently demonstrated by a randomly controlled trial of the anti-inflammatory glucocorticoid dexamethasone. Among hospitalized patients with COVID-19, the use of dexamethasone lowered 28-day mortality in patients receiving invasive mechanical ventilation (average age 59) and those receiving oxygen without ventilation (average age 66).^17^

This tremendous burden of infection in older adults is not limited to COVID-19. Between 12,000 to 61,000 deaths occur as a result of seasonal influenza in the United States each year, and approximately 90% of these deaths occur in patients ≥65 years of age.^18,19^ Between February and May 2020, the combination of deaths related to pneumonia, COVID-19, or seasonal influenza among all age groups accounted for 14% of deaths in the United States.^20^ Of these deaths, 9% were related to pneumonia, 7% to COVID-19, and 1% to influenza.^20^ Between 2009 and 2014, adults ≥60 years of age accounted for 70% of the sepsis cases reported in the United States.^21^ The mortality rate for sepsis was 16% for patients 60−79 years and 18% for patients ≥80 years of age.^21^ In another study, patients >80 years of age with sepsis had an in-hospital mortality rate of 47%.^22^

We have developed a system to discover and target the immune mechanisms that change with age and lead to dysregulated responses to infection. This system combines data generated from a high-throughput in vitro laboratory with machine learning to model the dysfunctional viral response in older adults. We trained the system to accurately detect the differences between young and old viral immune responses by exposing peripheral blood mononuclear cells (PBMCs) from young and old donors (N=89) to vesicular stomatitis virus encoding a red fluorescent protein (rVSV-ΔG-mCherry). This virus was chosen because of its ability to model the innate immune activation pathways of prevalent respiratory RNA viruses and because it is safe to use in a high-throughput, biosafety level one laboratory.^23,24^ The virus infected the monocytes and macrophages, which subsequently created a highly inflamed environment for the lymphocytes and other PBMCs. The differences between the young and old immune response were captured in our system’s multi-phenotype aging profile of viral response, which contained dozens of features measuring changes in cell composition, physical cell-to-cell interaction, organelle structure, cytokine levels, and other hidden complexities that contribute to age-related dysfunction. Many of these features showed significant differences between young and old viral responses. Since these aging profiles can be generated from a single well, we used them to assess the effect of 3428 bioactive compounds on the viral immune response of older adults (Figure 1). We ranked these compounds by combining the features in the multi-phenotype aging profiles into scores that measured their beneficial effect on immune aging and disease-specific phenotypes. Among others, the ranking revealed disulfiram and triptonide as two promising compounds that restored multiple aspects of the viral immune response of old PBMCs to a younger state. The scalability and generalizability of our system enables a novel characterization of the aging immune system that could be used to help sustain the health of older adults over their lifetime.

**Figure 1.**
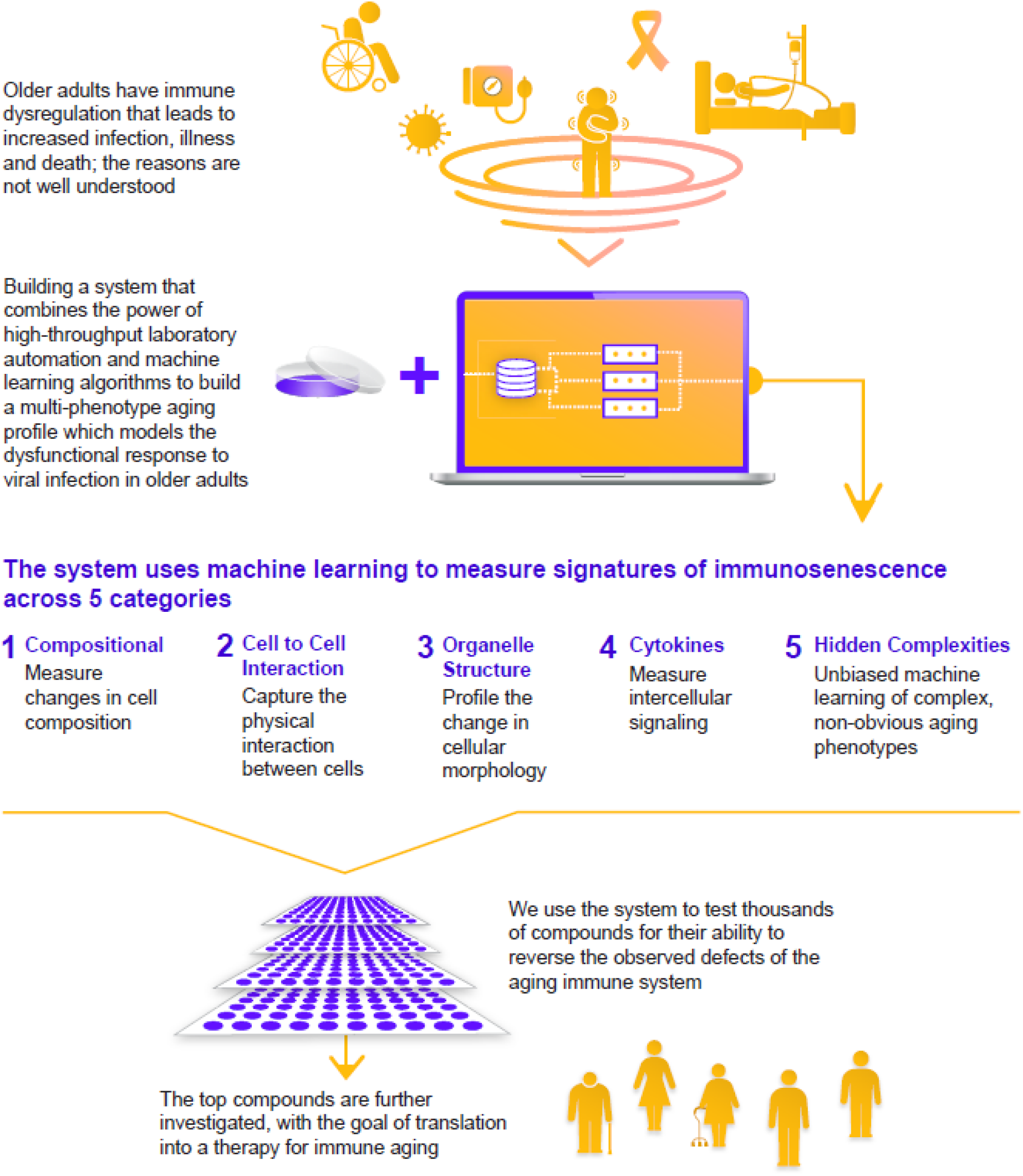
The system produces a multi-phenotype aging profile to measure immunosenescence.

## RESULTS

### Classification of Cell Types Based on Unbiased Morphological Analysis

The first feature we focused on in our multi-phenotype aging profile was cell composition. We built a scalable method for classifying immune cell types within bulk PBMCs based on cell morphology, because traditional methods for measuring cell composition were unable to scale to the needs of our high-throughput system and recent studies have shown that functional changes in cells can affect their shape while changes in shape can in turn alter their cellular function.^25–27^ To highlight a diverse range of morphological changes, the cells were stained with cell painting palettes that broadly characterized a variety of subcellular compartments such as the nucleus, cytoskeleton, mitochondria, and cell membrane.^28^ To understand and verify the morphological differences between immune cell types, we first isolated PBMCs from whole blood and then further isolated T cells (CD3^+^), B cells (CD19^+^), natural killer (NK) cells (CD56^+^), and monocytes (CD14^+^) using immunomagnetic negative selection based on customarily defined cell surface markers. Isolated cells were plated, stained with Cell Painting Palette 1 (Supplemental Table S1), and imaged. Within these images, we observed and quantified notable differences between the shapes of the isolated immune cell populations. As shown in Figure 2, the T cells and B cells shared the most similarities to each other; the T cells had a rounder and more uniform morphology, while the B cells had a more oval-shaped form with small areas of highly concentrated phalloidin staining on either end. The NK cells diverged from the tightly spherical form of other lymphocytes and showed more elongated and irregular forms. The monocyte populations were the largest in size and encompassed podosomes, dendrites, and kidney-shaped nuclei (Figure 2).

**Figure 2.**
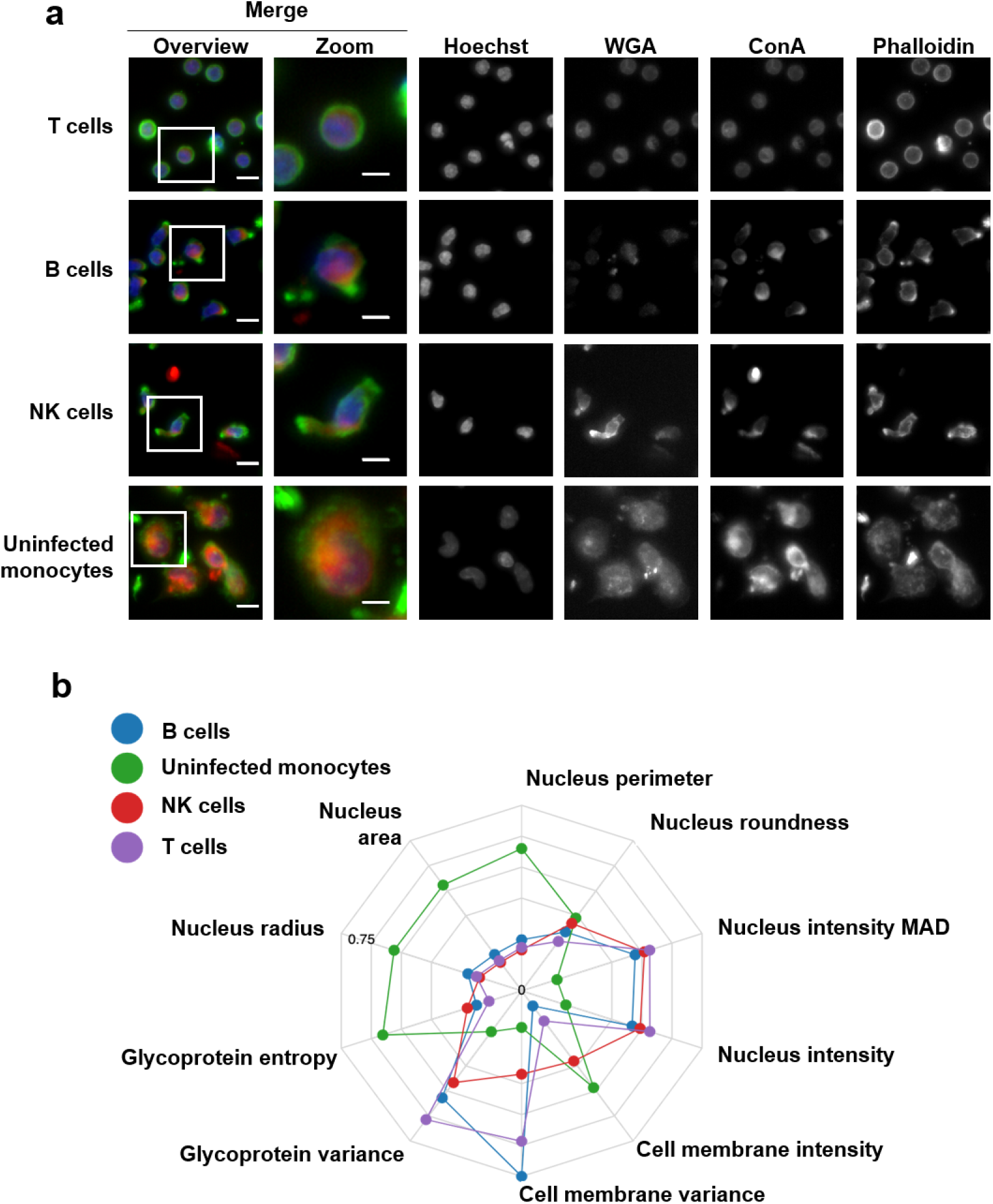
(a) Representative images of isolated T cells, B cells, NK cells, and monocytes from human bulk PBMCs using immunomagnetic negative selection and stained with Hoechst (blue), ConA (red), Phalloidin (green), WGA (far red). The boxed areas represent the single cell shown in the zoomed image. Scale bar = 10 μm (overview) and 5 μm (zoom). (b) Normalized averages for a subset of morphologic features that differentiate cell populations. ConA, concanavalin A; MAD, mean average deviation; NK, natural killer; PBMC, peripheral blood mononuclear cell; WGA, wheat germ agglutinin.

Using the observed phenotypic variation in isolated cells, we designed a computational model to classify single cells within a mixture of PBMCs. We built the model on PBMCs exposed to rVSV-ΔG-mCherry so that the multi-phenotype aging profile could accurately measure the compositional changes between rVSV-exposed PBMCs from young and old donors. These cells were stained with Cell Painting Palette 2, which revealed similar subcellular compartments as Cell Painting Palette 1, but included the membrane potential sensitive dye, MitoTracker Orange. The infected cells also expressed a mCherry-rVSV protein that had an overlapping emission spectrum with the MitoTracker causing them to be visualized in the same channel (Supplemental Table S2). Image embeddings, a statistically optimized representation of an image, were generated on an expert-curated training set of >61,000 single cell images from over 8 experimental runs using 31 plates seeded with cells from 105 distinct donors. The embeddings were then used to train a classification model. The model classified a single cell in an image of bulk PBMCs as a T cell, B cell, NK cell, dead cell and infected monocyte, or uninfected monocyte (Figure 3a). To validate the accuracy of the classification, we built an expert-labeled test set of >10,000 single cell images from PBMCs. When we evaluated the classifier on this data set, the model achieved 91% accuracy (data not shown). We also took single cell images and ran a principal component analysis on several hundred features derived from CellProfiler that were not used to train the model. Our model-predicted classification demonstrated clear clustering on the first two principal components (PC) of our analysis (Figure 3b). Because immune cell populations vary significantly between individuals, the model was run on PBMCs derived from several different individual donors to see if this variance could be captured by the classifier.^29^ Predicted cell compositions not only recapitulated the large inter-donor variance but also fell within the expected range of PBMC compositions (Figure 3c).^30^

**Figure 3.**
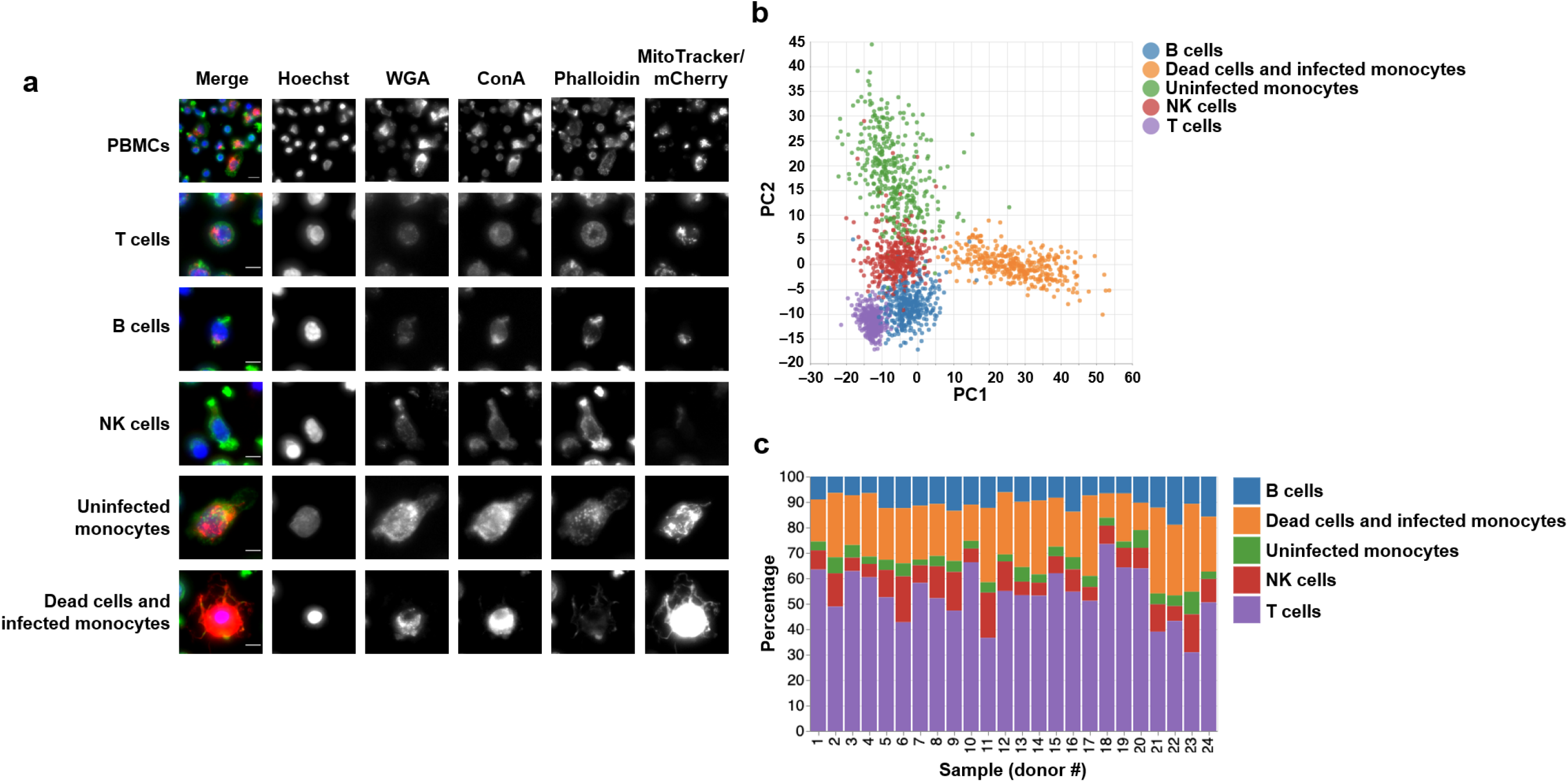
(a) Images of PBMCs and cells predicted to be T cells, B cells, NK cells, uninfected monocytes, or dead cells or infected monocytes. The cells were stained with the Cell Painting Palette 2 (Hoechst, blue; Phalloidin, green; MitoTracker/mCherry, red); WGA; ConA), and the predictions were derived from a model trained on image embeddings. Scale bar = 10 μm (PBMCs) or 5 μm (predicted cells). (b) PC analysis plot of 1200 morphologic features computed on single cells within bulk PBMCs exposed to rVSV-ΔG-mCherry. PCs were generated using the average feature value, then colored by cell type labels from a model trained on image embeddings of cell types. (c) The predicted PBMC composition of 24 donors exposed to rVSV-ΔG-mCherry at 10× MOI. The predictions were derived from a model trained on image embeddings computed from single-cell crops, labeled by cell type. ConA, concanavalin A; MOI, multiplicity of infection; NK, natural killer; PBMC, peripheral blood mononuclear cell; PC, principal component; rVSV-ΔG-mCherry, recombinant vesicular stomatitis virus expressing a red fluorescent construct; WGA, wheat germ agglutinin.

To provide an independent validation of our method, we correlated the cell composition predictions from our morphological model with those identified using flow cytometry (Supplemental Figure S1). The cell composition predictions from our model significantly correlated with those determined by flow cytometry for T cells (*r*=0.73; *P*=.007) and NK cells (*r*=0.92; *P*<.001). The dead cell and infected monocyte composition correlated significantly with all dead and infected myeloid cells derived from flow cytometry (*r*=0.62; *P*=.02; Figure 4). A B-cell marker was not included in the flow cytometry panel because of the mCherry overlap, so a correlation for the B-cell population was not calculated. While flow cytometry was able to accurately differentiate live and dead cells at different granularities, the morphological properties of dead cells were very similar to infected monocytes in the images, which made it difficult to differentiate live and dead infected cells using our system. Overall, the cell type classification model we developed provides a scalable and reproducible method to incorporate cell composition and other phenotypic features into our multi-phenotype aging profile.

**Figure 4.**
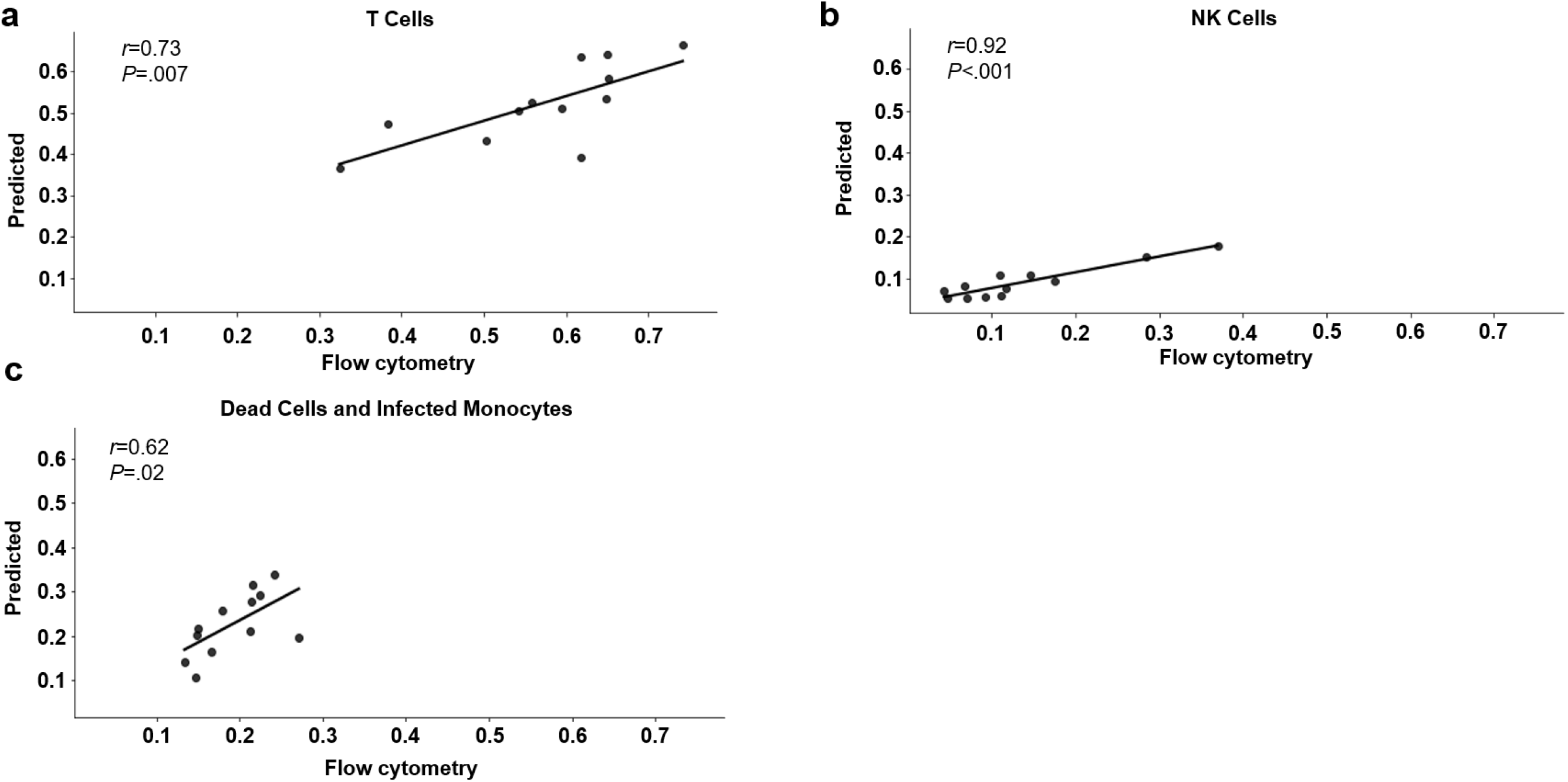
Correlation between the predicted composition of PBMCs exposed to rVSV-ΔG-mCherry and the composition determined by flow cytometry analysis for (a) T cells, (b) NK cells, and (c) infected and dead myeloid cells. Statistical analysis was performed via the 2-tailed *t* test. NK, natural killer; PBMC, peripheral blood mononuclear cell; rVSV-ΔG-mCherry, recombinant vesicular stomatitis virus expressing the fluorophore mCherry.

### Modeling the Inflammatory Antiviral Immune Response

Dysregulated immune responses are a central driver of disease progression in severe infection of older adults.^31–34^ To develop aging profile features that specifically measure phenotypes related to viral infection, we profiled the responses to viral load in PBMCs from donors ranging from 30 to 60 years old. We observed notable shifts in cellular composition in virally exposed PBMCs; specifically, there was a significant increase in dead cells and infected monocytes and a significant decrease in uninfected monocytes at 0.1×, 1×, and 10× multiplicity of infection (MOI; *P*<.001 for all MOIs; Figure 5a). The NK cell population decreased moderately but significantly for all viral conditions as compared to unexposed conditions (*P*<.001 for all MOIs; Figure 5a) but no difference was observed when comparing different viral loads. The T-cell and B-cell populations did not undergo significant changes due to viral load. We ran an experiment to test cytokine production of PBMCs when exposed to rVSV at 1×, 5×, 7× and 10× MOI. Secreted proinflammatory cytokines from the entire PBMC population increased in response to 10× viral exposure relative to 1× exposure, with significantly elevated levels of interleukin (IL)-6 (*P*<.001), tumor necrosis factor (TNF)α (*P*<.001), IL-1β (*P*<.001), and IL-8 (*P*<.001) at 10× MOI. In contrast, monocyte chemoattractant protein 1 (MCP1; *P*<.001) significantly decreased upon 10× viral exposure compared to 1× exposure (Figure 5b). The significant differences in these cytokines were also reinforced by similar dose dependent trends at 1×, 5×, 7×, and 10× MOIs (Figure 5b). Given the robust changes in PBMC composition and cytokine production upon viral exposure, we incorporated these features into our multi-phenotype aging profile to understand how they differ with age.

**Figure 5.**
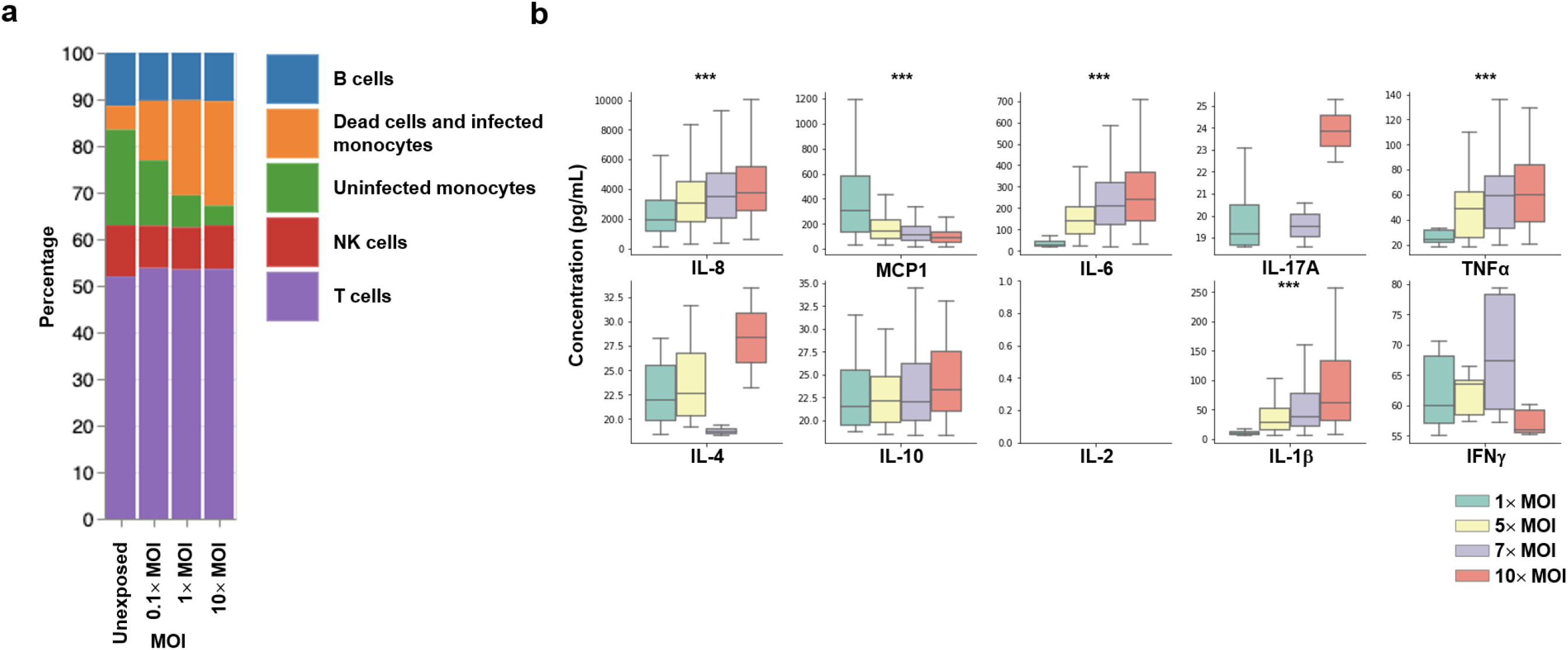
(a) Mean cell composition of 24 donors; cells were unexposed or exposed to rVSV-ΔG-mCherry at 0.1×, 1×, and 10× MOI for 24 h. (b) Cytokine levels in the supernatant 1×, 5×, 7×, and 10× rVSV-ΔG-mCherry-exposed PBMCs from 80 donors. The box and whisker plots indicate the median and quartiles. Statistical significance (via 2-tailed *t* test) is indicated; ***, *P*<.001. IFN, interferon; IL, interleukin; MCP1, monocyte chemoattractant protein 1; MOI, multiplicity of infection; NK, natural killer; PBMC, peripheral blood mononuclear cell; rVSV-ΔG-mCherry, recombinant vesicular stomatitis virus expressing a red fluorescent construct; TNF, tumor necrosis factor.

To more broadly measure how the immune response changes to different viral loads, we trained machine learning algorithms to learn the differences in the cellular immune response by titrating rVSV into PBMCs at different MOIs. We first generated embeddings on single cell images from PBMCs that were exposed to rVSV at 0.1×, 1×, or 10× MOI, and then stained with Cell Painting Palette 2. The embeddings were then used to train a machine learning classifier to discriminate the cellular responses to rVSV at a range of MOIs. PBMCs from 24 donors were evenly distributed across a 384-well plate assay to minimize bias due to plate location then exposed to rVSV at three different MOIs. We used stratified four-fold cross-validation to evaluate the classifier’s performance on each well. When we evaluated the model trained on monocytes only, the classifier was able to differentiate between unexposed, 0.1× MOI, 1× MOI, and 10× MOI with 91% accuracy across all four folds (Figure 6a). We trained a model on T cells but we were unable to distinguish between the unexposed cells and the cells exposed to 0.1× and 1× MOI conditions. However, our model learned significant differences between the T-cell response to 10× MOI cells and unexposed cells. The model was able to distinguish these conditions at an area under the receiver operating characteristic curve (AUC) of 0.92 across the four folds (Figure 6b). T cells, unlike monocytes, are not directly infected by the rVSV and therefore their response could be dependent on modifications of the environment after viral exposure which might explain why no difference is observed at lower MOIs. These validated models of cell response to viral loads can be used to observe whether the phenotype of old PBMCs looks more similar to the higher viral load phenotype compared with young PBMCs.

**Figure 6.**
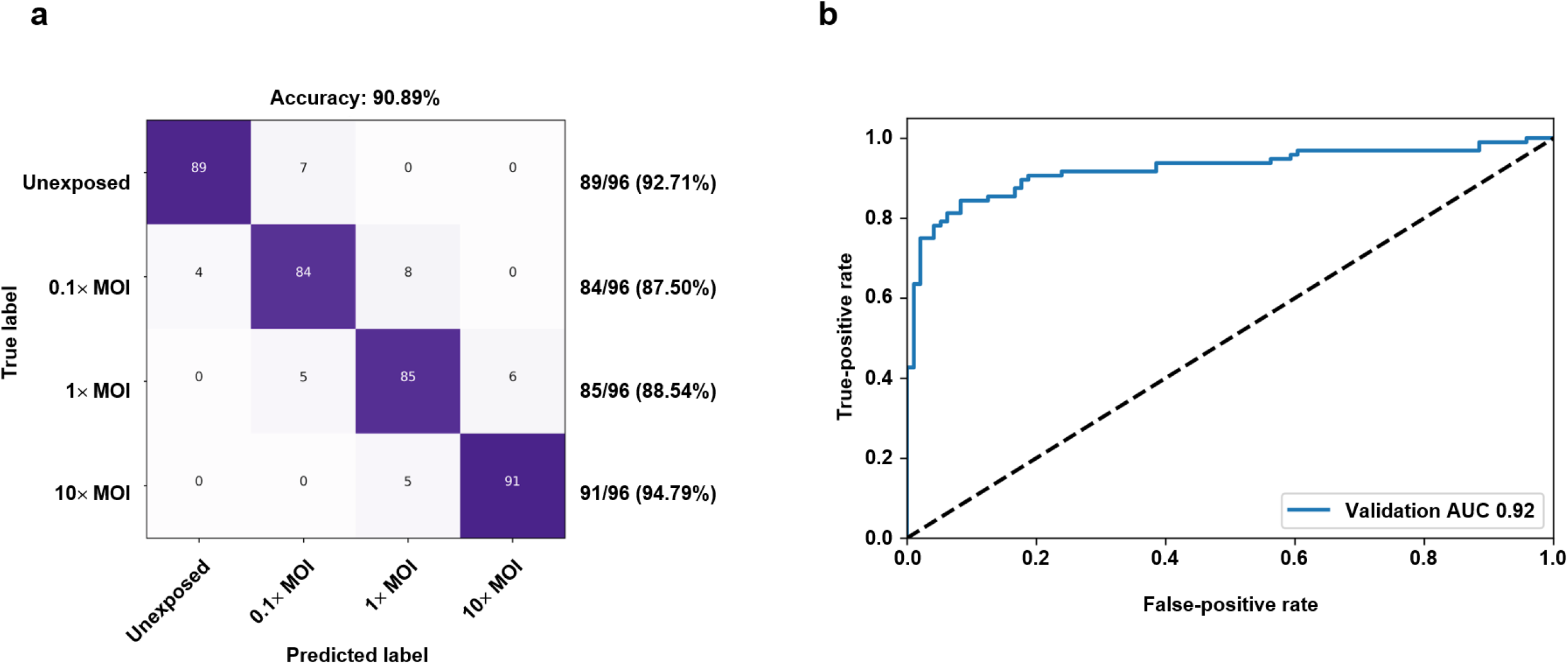
(a) Cross-validation performance of a single cell model trained on monocyte image embeddings to predict the different cellular phenotypes of no viral dose (unexposed), low viral dose (0.1× MOI), medium viral dose (1× MOI), and high viral dose (10× MOI) aggregated by taking the mean of fields in a well. Each condition represents 96 wells (4 replicates/MOI/donor). (b) Average cross-validation performance of a single cell model trained on T-cell image embeddings to predict the different cellular phenotypes of no viral dose (unexposed) and high viral dose (10× MOI). AUC, area under the receiver operating characteristic curve; MOI, multiplicity of infection.

### Dysfunction in the Viral Immune Response in Aging Adults

We used our multi-phenotype aging profile to conduct a novel characterization of the differences between young and old viral immune responses across dozens of features (Table 1). We generated the aging profiles for PBMCs from 49 young (<35 years of age) donors and 40 old (>60 years of age) donors. Demographics of the donors reflected the standard population of older adults, with old donors having a higher number of comorbidities, particularly hypertension, and taking more medications than young donors (Supplemental Table S3). All 89 donors were plated across two 384-well plate assays, with the two age populations evenly distributed across the plate to account for plate position effects. The donors were exposed to rVSV at 10× MOI and stained with Cell Painting Palette 2. Compared with young donors, PBMCs isolated from old donors had an increased percentage of dead cells and infected monocytes but showed no significant change in the percentage of uninfected monocytes (*P*<.001 and *P*=.53, respectively; Figure 7a). The percentage of T cells decreased in both age groups with a significant decrease in T cells from old donors compared with young donors (*P*<.001; Figure 7b). In contrast, a different experiment comparing unexposed PBMCs from young and old donors did not show a significant difference in T cells (*P*=.11). More work is necessary to understand these mechanisms, but the decreased T-cell population could recapitulate, at least in part, the lymphopenia observed in severely infected patients.^38,39^

**Figure 7.**
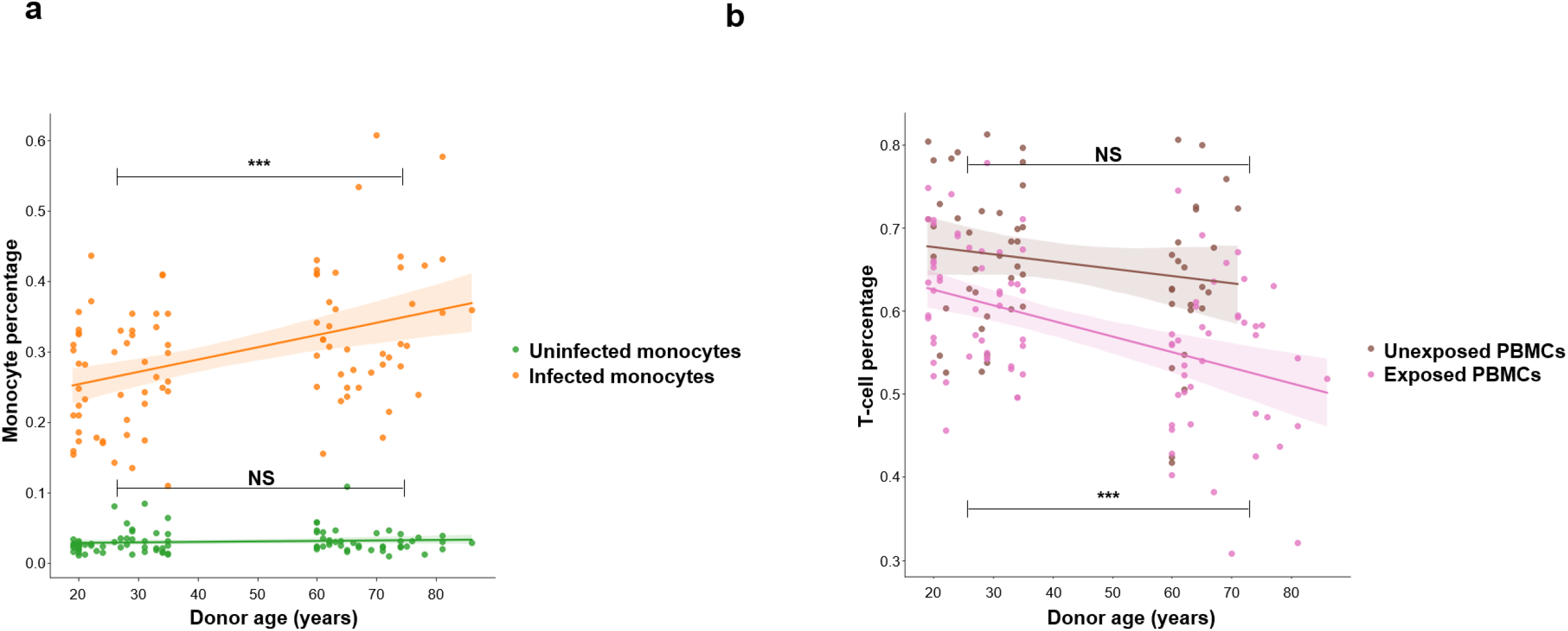
(a) Percentage of uninfected and rVSV-ΔG-mCherry-infected (at 10× MOI for 24 h) monocytes after viral exposure. (b) Percentages of T cells identified from unexposed and rVSV-ΔG-mCherry-exposed (at 10× MOI) PBMCs, presented according to donor age. Statistical significance (via two-tailed *t* test) between young and old donors after viral exposure is indicated (NS, nonsignificant; ***, *P*<.001). The shading around the trend lines indicates the 95% CIs of the fitted lines. MOI, multiplicity of infection; rVSV-ΔG-mCherry, recombinant vesicular stomatitis virus expressing a red fluorescent protein.

**Table 1.** Multi-Phenotype Aging Profile for Viral Response.

| Category | Metric |
| --- | --- |
| Compositional | % T cells<br>% Dead cells and infected monocytes<br>% Uninfected monocytes<br>% NK cells<br>% B cells |
| Cytokines | IFNγ, IL-1β, IL-2, IL-4, IL-6, IL-8, IL-10, IL-17A, MCP1, TNFα |
| Aging models | T cell<br>NK cell<br>Monocyte<br>PBMCs |
| Cell-to-cell interaction | Monocyte ↔ T cell<br>Monocyte ↔ Monocyte<br>T cell ↔ T cell |
| Structural | Reticular mitochondria<br>Concentric cytoskeleton<br>Mitochondrial texture |
| Viral load models | Monocyte viral load response<br>T-cell viral load response |

IFN, interferon; IL, interleukin; MCP1, monocyte chemoattractant protein 1; MOI, multiplicity of infection; NK, natural killer; PBMC, peripheral blood mononuclear cell; TNF, tumor necrosis factor.

Decreased T-cell levels in older individuals may be driven by cytokine-induced apoptosis or pyroptosis. This has been demonstrated to occur via induction of proinflammatory cytokines (eg, TNFα) and defects in the nuclear factor-κB (NF-κB) signaling cascade.^40,41^ To understand this relationship in aging, we compared the cytokine levels in our multi-phenotype aging profiles of young and old donors. No significant changes in cytokine production were observed between young and old donors in unexposed PBMCs (Figure 8a); there was also little difference in cytokine production of these cells when exposed to rVSV at 1× MOI (Supplemental Figure S2). However, when PBMCs were exposed to 10× MOI, significant differences in young and old cytokine production were observed. The production of proinflammatory cytokines significantly increased in PBMCs from old donors compared with those from young donors (IL-6 [*P*<.001]; MCP1 [*P*<.001]; TNFα [*P*<.001]; IL-1β [*P*<.001]; interferon [IFN]γ [*P*<.006]; IL-8 [*P*<.001]; Figure 8b) Importantly, these same cytokines have been implicated in the progression of hyperinflammation in diseases that disproportionally affect older adults, such as severe sepsis, COVID-19, and pneumonia. Interestingly, levels of IFNγ, TNFα, and IL-6 were also inversely correlated with the percentage of T cells in PBMCs exposed to rVSV at 10× MOI (IFN: old donors [*r*=–0.49; *P*=.02], young donors [*r*=–0.02; *P*=.93]; TNFα: old donors [*r*=–0.60; *P*<.001], young donors [*r*=–0.30; *P*<.09]; IL-6: old donors [*r*=–0.60; *P*<.001], young donors [*r*=–0.32; *P*<.02]; Figure 8c). Two recent studies of patients with COVID-19 demonstrated an inverse relationship between peripheral T-cell numbers and serum IL-6 and TNFα levels in patients over 60 years of age.^42,43^ The decrease in T-cell percentages and increase in proinflammatory cytokines that we observed in PBMCs obtained from old donors may provide a relevant in vitro model to study the progression of severe infection. A better understanding of the mechanisms driving these differences between young and old donors could therefore result in potential treatments for COVID-19 and reduce disease progression in older adults.

**Figure 8.**
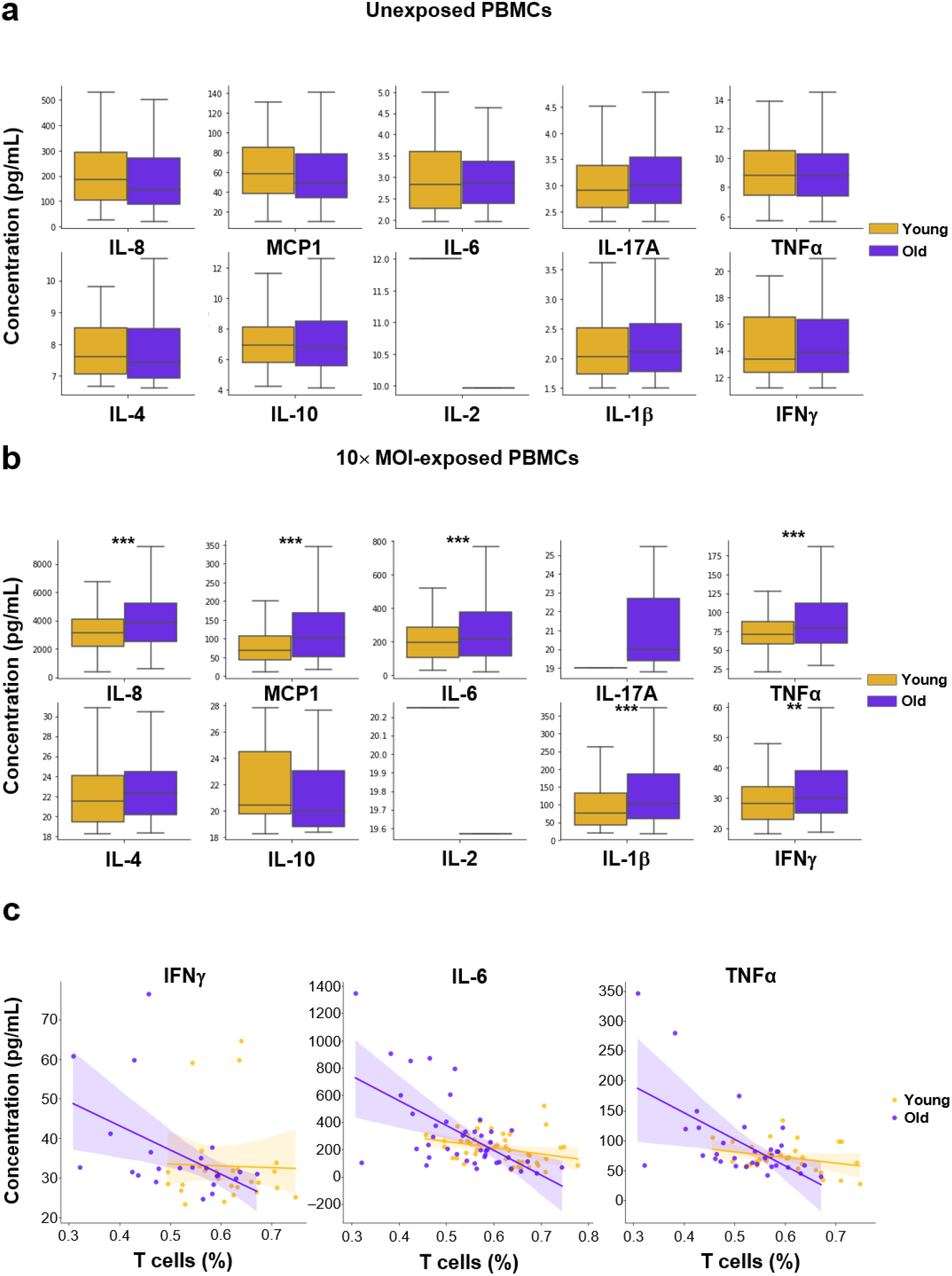
Cytokine levels from the supernatant of (a) unexposed and (b) rVSV-ΔG-mCherry-exposed (at 10× MOI for 24 h) PBMCs obtained from 89 donors from young and old adult populations. Statistical significance (via two-tailed *t* test) between young and old donors after viral exposure is indicated (**, *P*<.01, ***, *P*<.001). (c) Correlation between cytokine levels and percentage of T cells for PBMCs exposed to rVSV-ΔG-mCherry at 10× MOI. IFN, interferon; IL, interleukin; MCP1, monocyte chemoattractant protein 1; MOI, multiplicity of infection; PBMC, peripheral blood mononuclear cell; rVSV-ΔG-mCherry, recombinant vesicular stomatitis virus expressing a red fluorescent construct; TNF, tumor necrosis factor.

Next, we used image embeddings of T cells segmented from Cell Painting Palette 2-stained PBMCs to train an unbiased machine learning model to learn the hidden complexities that contribute to the differences between the viral immune response in young and old donors. We used stratified four-fold cross-validation to evaluate the classifier’s performance on each well. The model was able to differentiate between the antiviral T-cell response in cells obtained from young and old donors at an AUC of 0.90 (Figure 9a). The probability distribution of the test fold predictions was significantly different between young and old populations (*P*=.002). In several repeat validation experiments, the model was capable of significantly separating the same samples from young and old donors (*P*=.002; Figure 9b). This powerful tool can help discover modulators of previously unknown aging mechanisms.

**Figure 9.**
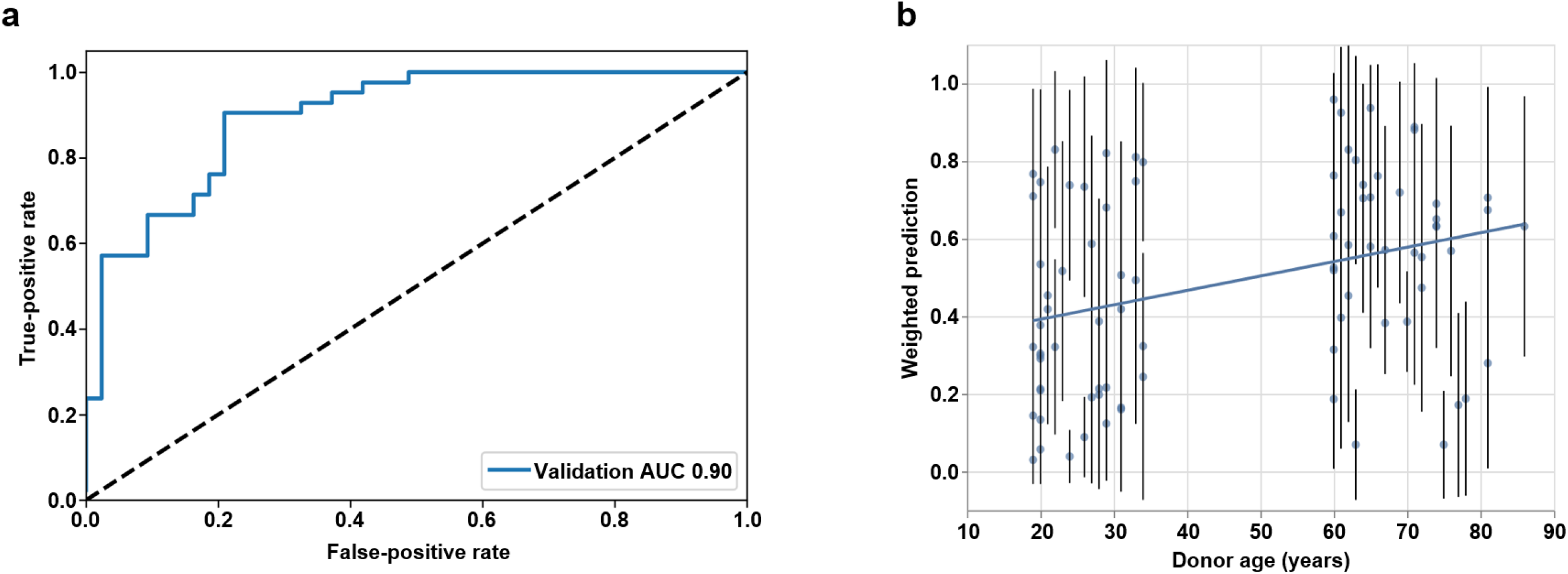
(a) Cross-validation performance of a model trained on T-cell image embeddings to predict the difference between immune responses in cells from young and old donors. (b) Weighted probability of samples being from old donors when the samples were obtained from a different experiment than the one used for training.

Cell composition, cytokines, and unbiased machine-learning–driven aging models are only a fraction of the features in our multi-phenotype aging profile. One big advantage of cell imaging is the ability to generate point patterns to model how interactions change with various perturbations.^37^ For example, interactions between T cells and antigen-presenting cells can lead to T-cell activation and aggregation of macrophages can contribute to cytokine production.^35,36^ To this end, we computed an interaction score using spatial statistical analysis to model non-Poisson multitype point distributions of specific cell types. The point patterns were assessed for complete spatial randomness using quadrate tests and distance methods. Spatial point pattern distributions of PBMCs were found to be homogenously distributed with strong evidence of interpoint interactions (Supplemental Figure S3). When we tested changes in cell-to-cell interaction between young and old donors, we found that the lymphocyte-to-monocyte interaction score was significantly increased in old donors compared with young donors (Figure 10a). The virus-related features in our aging profile also showed significant differences between young and old responses. Our T-cell viral load model predicts the T-cell response to the amount of viral load (0.1×, 1×, and 10× MOIs), and the same model predicted that the T-cell response of old donors had a higher apparent viral load than young donors (Figure 10a). Together, the many features within our multi-phenotype aging profile provide a comprehensive understanding of several differences between young and old donors.

**Figure 10.**
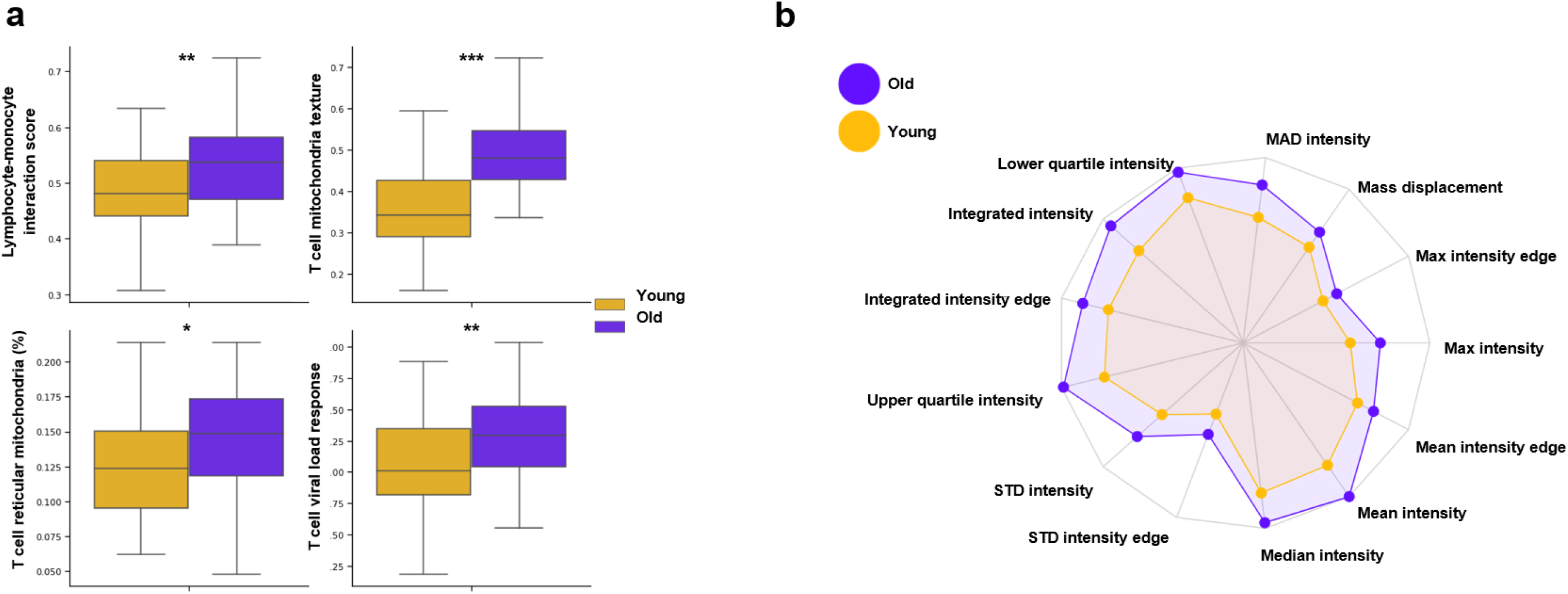
(a) Comparison of several features from our multi-phenotype aging profile between young and old donors. (b) Normalized averages for features measuring the intensity distribution of mitochondria membrane potential in T cells from old and young donors exposed to rVSV-ΔG-mCherry at a 10× MOI. AUC, area under the receiver operating characteristic curve; MAD, mitochondrial-associated protein degradation; MOI, multiplicity of infection. rVSV-ΔG-mCherry, recombinant vesicular stomatitis virus expressing a red fluorescent construct; STD, standard deviation.

In addition to the above, there are several features in the profile that provide a novel view of T-cell metabolism. The structural distribution of the mitochondrial network has been shown to be related to mitochondrial and cellular function. Spherical mitochondria, elongated mitochondria, and fused/reticular mitochondria all drive specific metabolic needs and functions.^44^ In our assay, we observed clear differences in the cellular distribution of mitochondria in T cells from young donors and old donors after viral exposure. The MitoTracker used in Cell Painting Palette 2 stains mitochondria in live cells and its accumulation is dependent upon membrane potential. The intensity of MitoTracker was significantly increased at the edge of the cell (*P*=.03) in old donors; we also observed a significant increase in the standard deviation of mitochondrial distribution throughout the cell (*P*=.003; Figure 10b). These measurements were complemented by specific features that calculate the percentage of reticular mitochondria and changes in the overall texture of the mitochondria. The percentage of reticular mitochondria was significantly increased in old adults compared with young adults, while a model of mitochondrial texture also showed significant differences (*P*=.04; *P*<.001; Figure 10a). These general changes in mitochondrial structure between young and old cells are consistent with reported mitochondrial mechanisms that have been causally linked to the age-related T-cell senescence and decline in T-cell function.^45^ Compounds that alter the mitochondrial behavior in cells from old donors, or any of the other signatures we modeled in our system, may improve the response to infection in older adults.^46^ Altogether, there are many significant differences between young and old viral immune responses as measured by dozens of features in our multi-phenotype aging profile.

### Evaluating the Multi-Phenotype Aging Profile of 3428 Bioactive Compounds

To find promising therapeutics for the purpose of rejuvenating immune responses in older adults, we generated our multi-phenotype aging profile for each of 3428 bioactive compounds. Of the 3428 compounds, approximately 1900 have been approved and marketed in at least one country. PBMCs from 35 young and old donors were seeded in 384-well plates and exposed to rVSV, then the PBMCs from old donors were treated with a single compound or vehicle to be used as controls. Each individual treatment and control well generated its own multi-phenotype aging profile containing dozens of age-related features. Every feature in the aging profile was classified as a “hit” or “miss” based on the magnitude of its distance from old vehicle-only controls. All labeled features were combined into scores for the aging profile of each compound by weighting each feature based on its relevance to immune aging or COVID-19. For our COVID-19 cytokine score, we increased the weights of cytokines reported to be increased in patients with severe COVID-19 so that compounds moving these cytokine levels in a beneficial direction ranked more highly. For our immune aging score, we weighted the features based on the consistency and magnitude of the differences between young and old donors. Generally, this method allows us to tailor scoring in order to focus on any immune dysfunction in any specific disease by altering weights of the features. For example, a disease like cancer could have a score that more heavily weighs the cell-to-cell interaction features to find potential therapeutics that improve T-cell trafficking.

These immune aging and COVID-19 cytokine scores allowed us to evaluate thousands of compounds for their therapeutic potential by measuring how each one affects dozens of age-related phenotypes in rVSV-exposed PBMCs. All compounds were ranked by our COVID-19 cytokine and immune aging scores, then the top 196 highest-scoring compounds were validated in a follow-up dose-response study. Through this process, we identified two compounds, disulfiram and triptonide, with the potential for rejuvenating the aging immune system (Figure 11). We are sharing two of the top ten hits that are exemplary of our results. We chose to share our disulfiram results in particular as we believe it is a cheap and widely available drug with a promising safety profile that may be immediately therapeutically useful.

**Figure 11.**
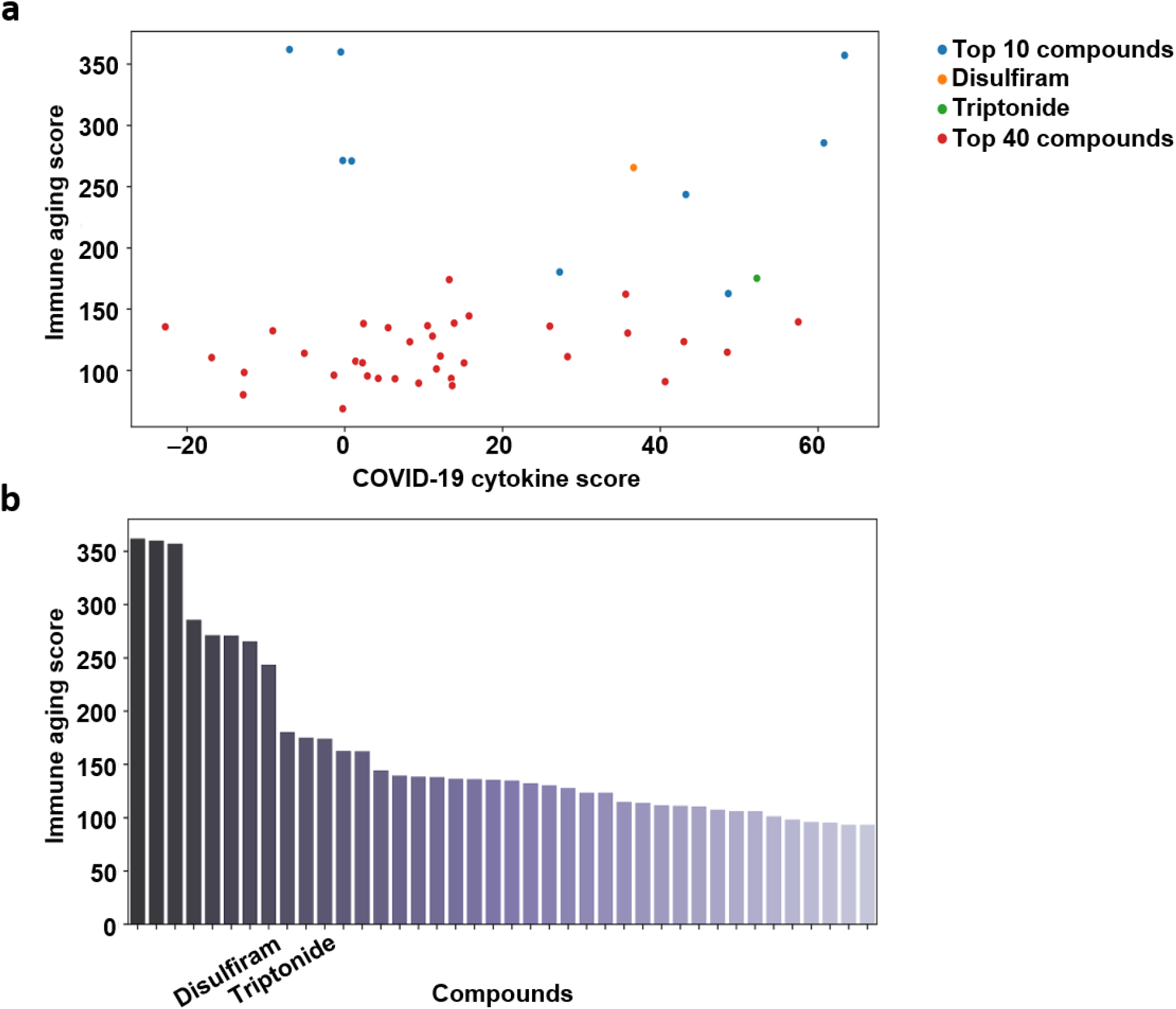
(a) Immune aging score and COVID-19 cytokine score plotted for the top 40 compounds from our high throughput screen of 3428 bioactive compounds. Ranking defined by the sum of the immune aging score and the COVID-19 cytokine score. (b) A ranking of our immune aging score for our top 40 compounds. COVID-19, coronavirus disease 2019.

### Evaluation of Triptonide’s Effect on the Multi-Phenotype Aging Profile of Old PBMCs

Triptonide, a diterpenoid from the medicinal herb *Tripterygium wilfordii* Hook F,^47^ had notable effects on virally exposed PBMCs from old donors. *Tripterygium* compounds generally have demonstrated anti-inflammatory, immunosuppressive, and antiviral effects in animal models and in vitro studies.^47–49^ Studies have shown that the anti-inflammatory effects of *Tripterygium* compounds target both T cells and macrophages. For T cells, activation is suppressed through inhibition of NF-κB transcription.^50^ For macrophages, IL-1β, IL-6, and TNFα production are decreased and expression of the anti-inflammatory cytokine IL-37 is increased.^51,52^

Consistently, we observed significant decreases in proinflammatory cytokines produced by triptonide-treated rVSV-exposed PBMCs relative to old vehicle-only control PBMCs. After normalizing the cytokine levels by the total number of cells, MCP1, IL-1β, and TNFα all showed significant differences at one or more concentrations tested compared with vehicle-only controls (Figure 12). The most notable of these was MCP1, which showed a significant, dose-dependent reduction when compared with vehicle-only controls (Figure 12). These results confirm what was previously seen in the literature for the *Tripterygium* compounds and validate the ability of our aging profile to identify potential immunomodulatory compounds.^47,48^ Triptonide also shifted the T-cell metabolic features from an older profile to a younger profile. There was a concentration-dependent decrease in the percentage of reticular T-cell mitochondria (Figure 13a), consistent with reports that indicate that activated T cells have more fragmented mitochondrial networks, and activated T cells may be more prevalent in responsive, relatively young immune systems.^53^

**Figure 12.**
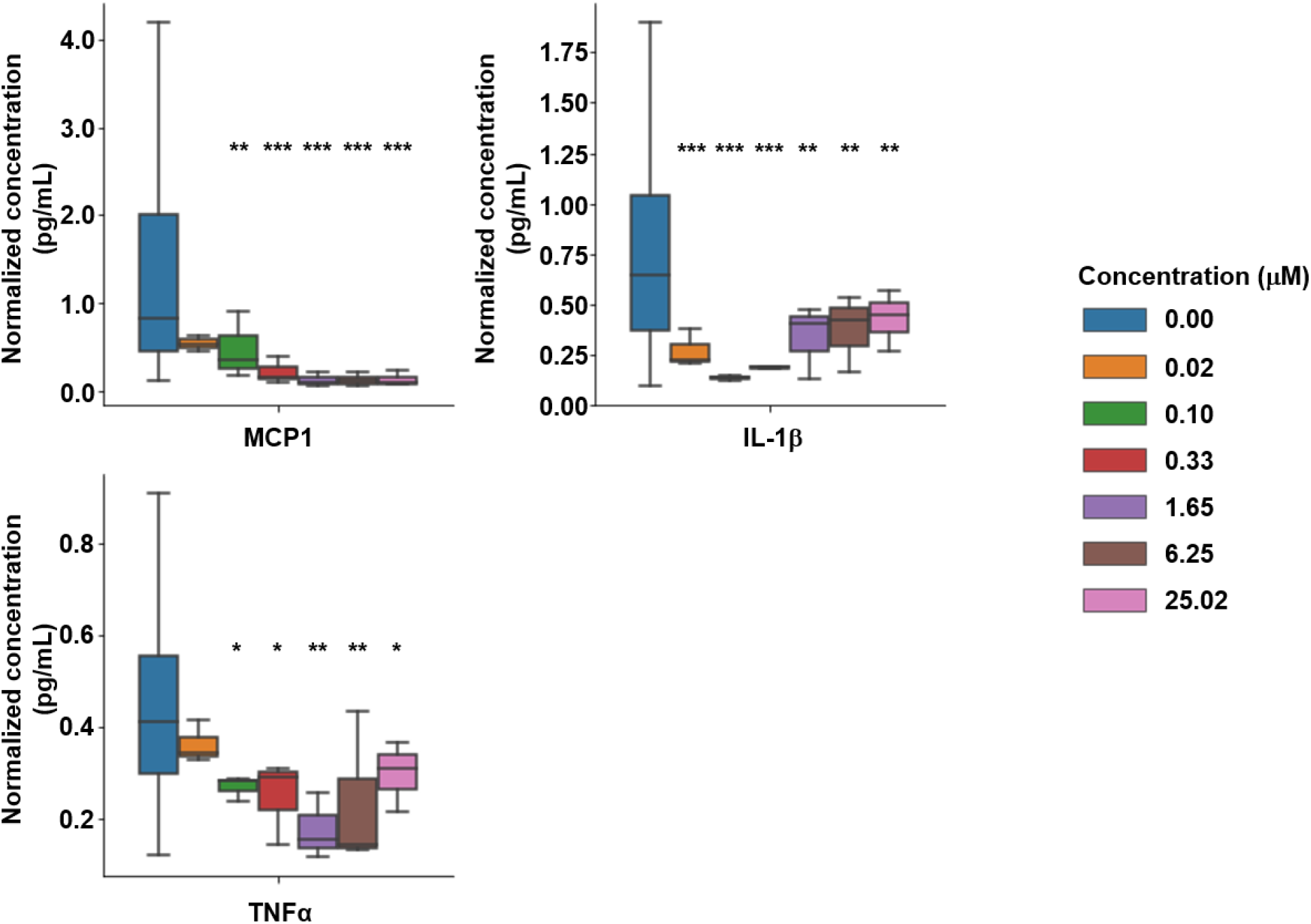
Cytokine levels from the supernatant of rVSV-ΔG-mCherry-exposed (at 10× MOI) PBMCs from three old donors and treated with different concentrations of triptonide. The *P* values for each donor were combined using the Fisher’s combined probability method. Asterisks indicate statistical significance (via two-tailed *t* test) relative to vehicle-only control cells: \**P*<.05; \*\**P*<.001; \*\*\**P*<.0001. IL, interleukin; MCP1, monocyte chemoattractant protein 1; MOI, multiplicity of infection; PBMC, peripheral blood mononuclear cell; TNF, tumor necrosis factor.

**Figure 13.**
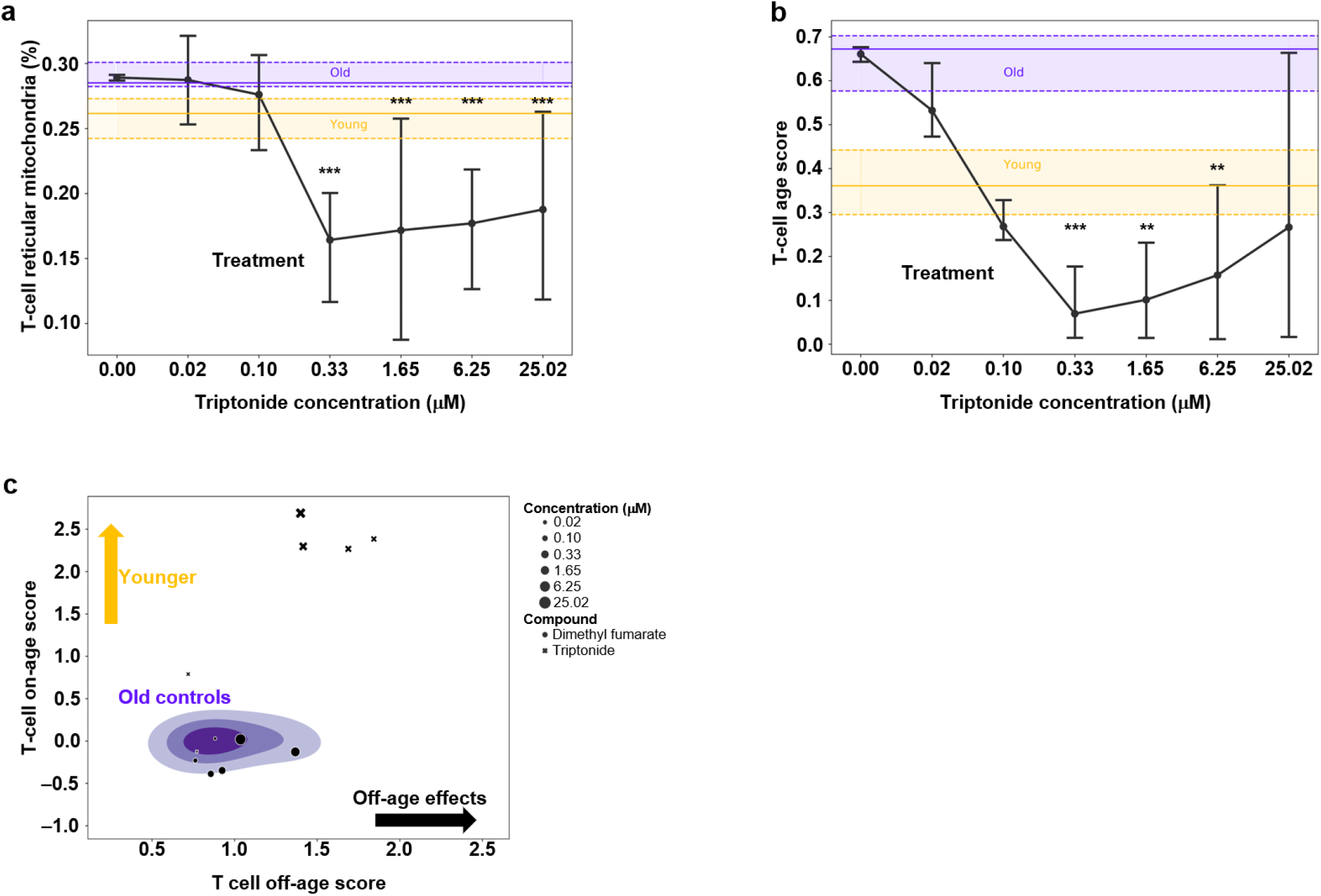
PBMCs were obtained from three old donors and treated with different concentrations of triptonide. The *P* values for each donor were combined using the Fisher’s combined probability method. (a) Reticularity measurement of mitochondria (b) and aging scores in T cells. Horizontal bars represent the distribution of vehicle-only controls from young (yellow) and old (purple) donors; the solid line represents the median and the lower and upper dashed lines represent the 25th and 75th quartiles, respectively. The line graph represents the median and 95% CI for treated cells, with statistical significance relative to vehicle-only control cells indicated by asterisks. (c) On-age and off-age scores for T cells treated with either triptonide or dimethyl fumarate. Distributions for young and old control (vehicle only) cells are plotted using a Gaussian kernel density estimation. Statistical significance (via two-tailed *t* test) relative to vehicle-only control cells is indicated by the following: \**P*<.05; \*\**P*<.001; \*\*\**P*<.0001.

The immunosuppressive and anti-inflammatory effects of *Tripterygium* compounds have also been linked to their ability to induce apoptosis in T cells.^54^ We saw that triponide-treated PBMCs had a significant increase in the percentage of dead cells and infected monocytes and a decrease in the percentage of T cells when compared to vehicle-only controls. For the highest three concentrations, there was an average of 21% more dead and infected monocytes and 19% less T cells relative to vehicle-only controls (data not shown). This was a common trend for many of our top scoring compounds.

The predictions from our T-cell aging models also significantly changed after treatment with triptonide. Compared with old vehicle-treated controls, PBMCs from old donors treated with various doses of triptonide shifted the model’s readout from an older phenotype to a younger phenotype (Figure 13b). At a low concentration, the T-cell age score of triptonide-treated PBMCs did not significantly change relative to old vehicle-only controls but significantly decreased with treatment concentrations above 0.33 μM (0.02 μM [*P*=.85]; 0.1 μM [*P*=.16]; 0.33 μM [*P*<.001]; 1.65 μM [*P*=.001]; 6.25 μM [*P*=.008]; 25.02 μM [*P*=.005]; Figure 13b). To better understand this shift in T-cell age score after treatment, we used image embeddings of T cells to compute an “on-age” score, measuring the distance from young controls, and a complementary “off-age” score, measuring the magnitude of the change in the orthogonal direction to age distances. Several concentrations of triptonide significantly shifted the “on-age” score closer to vehicle-only cells from young donors and farther from that obtained using vehicle-only cells from old donors (0.02 μM [*P*=.27]; 0.1 μM [*P*<.001]; 0.33 μM [*P*<.001]; 1.65 μM [*P*<.001]; 6.25 μM [*P*<.001]; 25.02 μM [*P*<.001]; Figure 13c). Triptonide also induced significant “off-age” effects (0.02 μM [*P*=.007]; 0.1 μM [*P*=.43]; 0.33 μM [*P*<.001]; 1.65 μM [*P*<.001]; 6.25 μM [*P*<.001]; 25.02 μM [*P*<.001]; Figure 13c). These “off-age” effects are not definitively detrimental but show that the compound induces effects that are not directly related to age. In our compound screen, we observed all possible effect combinations; some compounds only showed “on-age” effects, some showed both “on-age” and “off-age” effects, some showed neither, and some induced only “off-age” effects. As an example, another compound, dimethyl fumarate, did not improve the “on-age” effects, as indicated by “on-age” scores that were very close to those of old vehicle-only controls (Figure 13c).

Taken together, the multi-phenotype aging profile of old rVSV-exposed PBMCs treated with triptonide showed restored phenotypes in cytokines, reticular mitochondria, and the T-cell predicted age at multiple concentrations. A deeper understanding of the aging mechanisms behind triptonide could shed light into the dysfunction of immune viral responses in older adults.

### Evaluation of Disulfiram’s Effects on the Multi-Phenotype Aging Profile of Old PBMCs

In addition to triptonide, the small molecule disulfiram also demonstrated significant beneficial effects on our multi-phenotype aging profile of PBMCs from old donors exposed to rVSV. Disulfiram is an approved alcoholism aversion therapy that blocks the reaction converting acetaldehyde, an alcohol metabolism intermediate, into acetate.^55,56^ These effects are complemented by several anti-inflammatory mechanisms, including an ability to inhibit inflammasome-mediated pyroptotic cell death.^57,58^

Within our aging profiles, disulfiram significantly reduced several proinflammatory cytokines when compared with old vehicle-only controls from old donors. After normalizing the cytokine levels by the total number of cells, MCP1, IL-1β, IL-6, and TNFα were all significantly decreased for at least two concentrations (Figure 14). The reduction in MCP1 was concentration-dependent (Figure 14). Similarly, IL-1β was significantly reduced at 0.1 μM, and this effect was maintained for all subsequently higher concentrations of disulfiram (Figure 14). These anti-inflammatory effects rank disulfiram highly in our COVID-19 cytokine score, showing its potential to improve older viral immune responses to infection.

**Figure 14.**
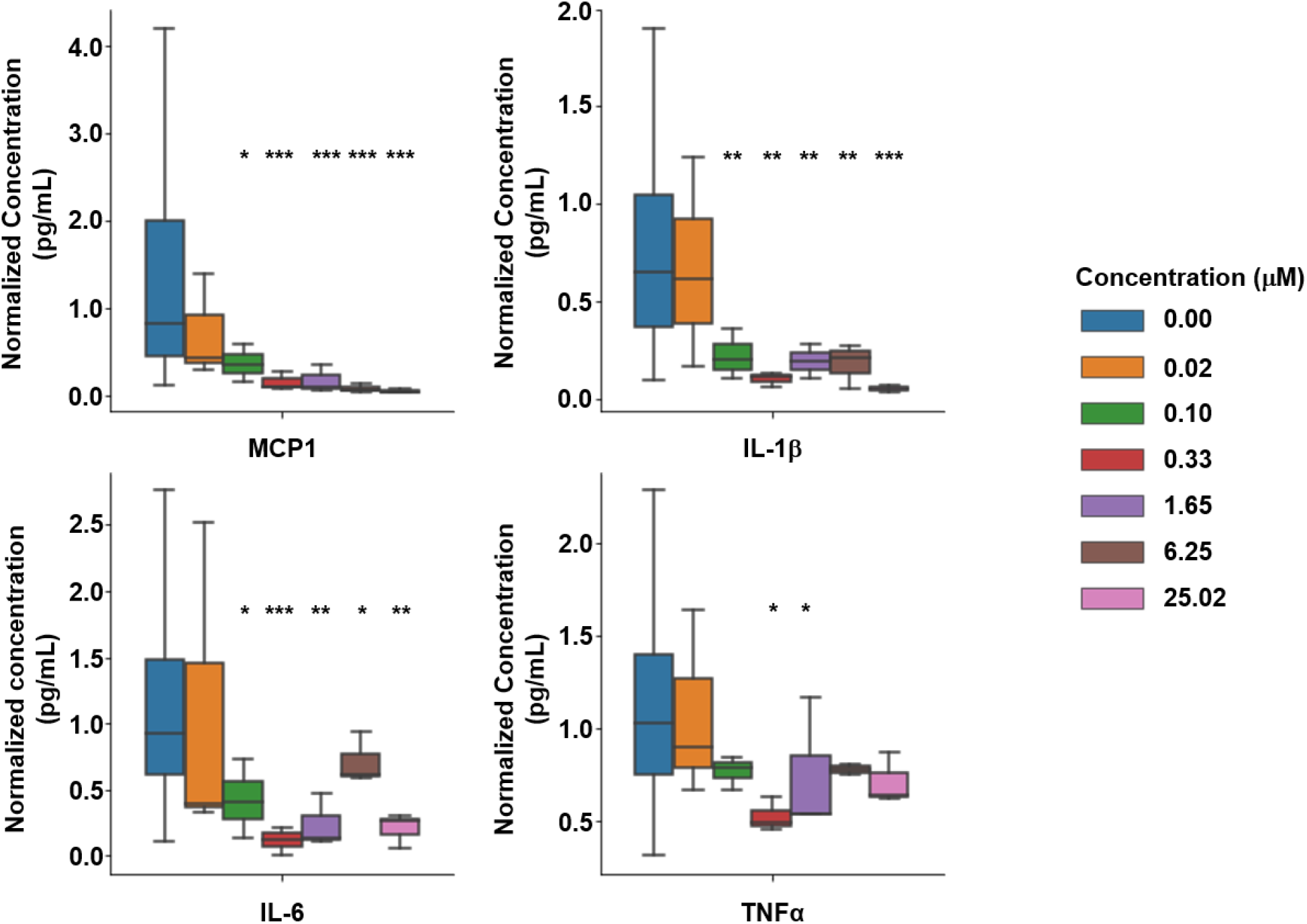
Cytokine levels from the supernatant of rVSV-ΔG-mCherry-exposed (at 10× MOI) PBMCs obtained from three old donors and treated with different concentrations of disulfiram. Asterisks indicate statistical significance (via two-tailed *t* test) relative to vehicle-only control cells: \**P*<.05; \*\**P*<.001; \*\*\**P*<.0001. IL, interleukin; MCP1, monocyte chemoattractant protein 1; MOI, multiplicity of infection; PBMC, peripheral blood mononuclear cell; TNF, tumor necrosis factor.

Disulfiram treatment induced similar compositional changes as the other top compounds. On average, the three highest concentrations of disulfiram increased dead cells and infected monocytes by 12% and decreased T cells by 13%. This trend is in the opposite direction of a young phenotype but consistent with disulfiram’s ability to induce apoptosis through redox-related processes.^59^

The viral features within our multi-phenotype aging profile were also significantly improved after treatment with disulfiram. Vehicle- and disulfiram-treated PBMCs were exposed to 10× MOI rVSV, and at several concentrations, disulfiram caused the cells to appear like they were responding to a lower viral load. In particular, compared with old, vehicle-only controls, concentrations of disulfiram above 1.65 μM consistently and significantly made the cells appear to be responding to a lower viral load (0.02 μM [*P*=.63]; 0.1 μM [*P*=.01]; 0.33 μM [*P*=.43]; 1.65 μM [*P*=.05]; 6.25 μM [*P*<.001]; 25.02 μM [*P*<.001]). Additionally, disulfiram treatment exceeding 0.33 μM also significantly reduced the percentage of T cells displaying reticular mitochondrial networks (0.02 μM [*P*=.04]; 0.1 μM [*P*=.42]; 0.33 μM [*P*<.001]; 1.65 μM [*P*<.001]; 6.25 μM [*P*<.001]; 25.02 μM [*P*<.001]; Figure 15a). Importantly, these phenotypes restored by disulfiram are both found at higher viral loads and are significantly different between young and old viral immune responses.

**Figure 15.**
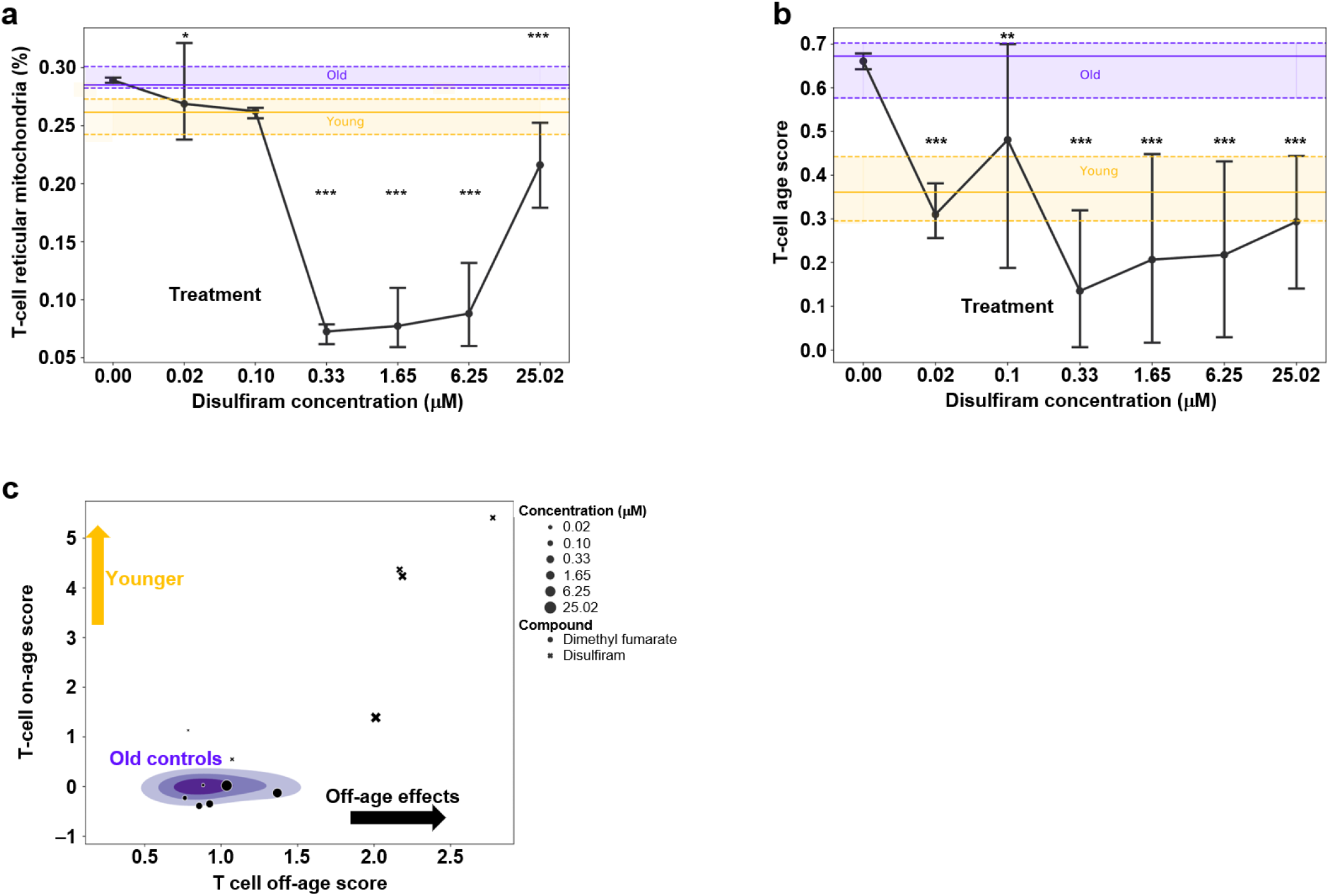
PBMCs were obtained from three old donors and treated with different concentrations of disulfiram. The *P* values for each donor were combined using the Fisher’s combined probability method. (a) Reticularity measurement of mitochondria (b) and aging scores in T cells. Horizontal bars represent the distribution of vehicle-only controls from young (yellow) and old (purple) donors; the solid line represents the median and the lower and upper dashed lines represent the 25th and 75th quartiles, respectively. The line graph represents the median and 95% CI for treated cells, with statistical significance relative to vehicle-only control cells indicated by asterisks. (c) On-age and off-age scores for T cells treated with either disulfiram or dimethyl fumarate. Distributions for young and old control (vehicle only) cells are plotted using a Gaussian kernel density estimation. Statistical significance (via two-tailed *t* test) relative to vehicle-only control cells is indicated by the following: \**P*<.05; \*\**P*<.001; \*\*\**P*<.0001.

T cells from rVSV-exposed PBMCs treated with disulfiram also exhibited a significantly younger phenotype compared with old vehicle-only controls. Disulfiram showed a significant rejuvenating effect on the T-cell age scores at all concentrations in our multi-phenotype aging profiles (0.02 μM [*P*<.001]; 0.1 μM [*P*=.005]; 0.33 μM [*P*<.001]; 1.65 μM [*P*<.001]; 6.25 μM [*P*<.001]; 25.02 μM [*P*<.001]; Figure 15b). Similar trends were seen with the “on-age” and “off-age” scores. Compared with vehicle-only controls, the “on-age” T-cell score significantly shifted in the young direction after treatment with disulfiram (0.02 μM [*P*=.003]; 0.1 μM [*P*=.11]; 0.33 μM [*P*<.001]; 1.65 μM [*P*<.001]; 6.25 μM [*P*<.001]; 25.02 μM [*P*<.001]; Figure 15c). The “off-age” score also had a significant effect in the samples exposed to concentrations of disulfiram above 0.33 μM (0.02 μM [*P*=.26]; 0.1 μM [*P*=.44]; 0.33 μM [*P*<.001]; 1.65 μM [*P*=.008]; 6.25 μM [*P*=.003]; 25.02 μM [*P*=.01]; Figure 15c).

Altogether, these beneficial shifts in age-related phenotypes demonstrate the strong potential of disulfiram for treating dysfunction in the old viral immune response.

## DISCUSSION

Aging is the most critical risk factor for morbidity, mortality, and progression of severe infectious disease.^21,60–62^ While the COVID-19 pandemic has drawn attention to the vulnerability of older adults, the heightened risk faced by these patients is not limited to this pandemic.^63^ Older adults face an increased risk of death from seasonal influenza and sepsis, and the ever-increasing aging population across the globe presents a challenge to protect this at-risk population. To help address this challenge, we developed a system that produces a multi-phenotype aging profile by combining high-throughput lab automation and machine learning algorithms. This is a novel way to study immune cell responses to viral infection that allows for the rapid characterization of these responses in multiple dimensions (eg, cell composition, cell-to-cell interactions, cellular features, cytokine production, and hidden complexities) and enables us to screen for compounds that can have an impact on how immune systems of older patients respond to viral infections.

There were some limitations to our approach. First, we could not differentiate between dead cells and those infected with rVSV given their morphological similarities. To resolve this, we grouped these cells into one class called “dead cells and infected monocytes.” Although we were still able to identify multiple significant effects, future work will focus on building more granular cell types that differentiate between infected cells and cells that died from apoptosis, pyroptosis, or other cell death pathways. In addition, the presence of dead cells could have impacted the compositional readouts. Since our in vitro system cannot naturally add or replenish cells, most changes in composition are driven by dying and nonadherent cells rather than cell trafficking, proliferation, or differentiation. Additionally, Cell Painting Palette 2 had significant overlapping fluorescent signals in the wavelengths between 550nm–610nm. We chose to combine MitoTracker Orange (554/576 ex/em) and mCherry (578/610 ex/em) into a single panel given the lack of infection in cell types other than monocytes. While mCherry generated a distinct morphological phenotype upon expression after infection, the strength of this mCherry signal covered the MitoTracker signal in alive but infected cells.

Another limitation is the potential for confounders to unknowingly bias our machine learning models. A significant amount of our top compounds saw a compositional shift which was in the opposite direction of a young phenotype. For many of these compounds, dead cells and infected monocytes significantly increased while T cells significantly decreased. We hypothesized that this increase was mostly due to cells undergoing compound induced apoptosis then being labeled as dead cells and infected monocytes. While our T-cell machine learning aging models were not trained or evaluated on cells from this class, the ability of these models to capture subtle trends from high dimensional data leaves them vulnerable to bias by technical effects. This requires special tooling and attention to ensure known confounders, such as plate, donor, and batch effects, are not biasing the models and that proper awareness is brought to new potential confounders. The many features in our multi-phenotype aging profile allow us to broadly profile compound effects leading to the identification of some features, such as apoptosis, which could be potential confounders. We do not know if an increase in apoptosis impacted the model or whether the impact would be a negative artifact rather than a real beneficial aging phenotype. One hypothesis is that the increase in apoptosis caused differential cytokine release which induced an early stimulation of the innate immune system so that it was more prepared to mount a better viral response and resolution.^64^ More work is needed to test these results for specific mechanistic hypotheses and in vivo efficacy.

Despite these limitations, we conducted a successful screen on 3428 bioactive compounds. Through this screen, we identified leading candidates for treatment of severe infection by assigning scores to many features within our aging profile based on their relevance to immune aging and COVID-19 cytokine pathology. Within the top 10 ranked compounds from our screen, disulfiram, a long-used therapy for patients with alcohol dependency, and the diterpenoid triptonide, obtained from a woody vine commonly found in Eastern and Southern China, were two interesting compounds that robustly rejuvenated aspects of the viral immune response in old rVSV-exposed PBMCs. The reported anti-inflammatory mechanisms of triptonide in combination with findings from our study indicate that this compound shows promise for dysfunctional aging immune responses. However, additional research is needed to ensure its safety before use in humans for aging-related indications. *Tripterygium* compounds have demonstrated liver and genital toxicity in animal models.^65^ While the toxicity of triptonide may make it unsuitable for the clinic, further understanding of its mechanisms could inform a search for similar compounds that would be more suitable for human therapeutic use. Further studies are in progress to better understand the mechanisms by which triptonide modulates immune cells in the hope to identify specific compounds that target similar age-related mechanisms.

In contrast to triptonide, disulfiram has over 60 years of clinical use in humans. It has been used as an alcohol aversion therapy through its inhibition of aldehyde dehydrogenase.^56,66^ Over these 6 decades, disulfiram has an established and well-understood safety profile with no significant evidence of clinical toxicity in the absence of ethanol.^66^ Our multi-phenotype aging profile of disulfiram demonstrated significant anti-inflammatory and rejuvenating effects. It reduced the proinflammatory cytokines MCP1, IL-1β, IL-6, and TNFα, and rejuvenated several other features in our aging profile such as T-cell age score, viral load response, and mitochondrial morphology. These mechanisms could be related to the recently discovered ability of disulfiram to block the final step in inflammasome–mediated pyroptosis and cytokine release.^67^ Disulfiram blocks gasdermin D (GSDMD), a pore-forming protein that plays a critical role in the terminal step of inflammasome-mediated pathways by facilitating the release of proinflammatory cytokines.^68–70^ Cleaved GSDMD similarly creates pores on the cell membrane of neutrophils to allow for the release of neutrophil extracellular traps (NETs), a process known as NETosis.^71,72^ Both of these mechanisms make disulfiram an attractive treatment for hyperinflammatory infections such as COVID-19, Acute Respiratory Disease Syndrome (ARDS), and sepsis.^58,73,74^ Disulfiram has already been shown to protect mice from lethal lipopolysaccharide-induced septic shock.^67^ There is mounting evidence that targeting NETosis and inflammasome activation could be a potential treatment for severe COVID-19. Markers in the sera of patients with severe COVID-19 indicate the presence of NETs, and an autopsy of a lung specimen from a patient with COVID-19 showed extensive neutrophil infiltration.^75,76^ In addition, proinflammatory cytokines in patients with severe COVID-19 were significantly higher than in moderate cases.^77^ This includes elevated levels of IL-1β which is typically a result of inflammasome activation.^78^ Given the evidence of NETosis and inflammasome-mediated pyroptosis in severe COVID-19 pathogenesis, blocking the terminal step of both of these pathways by inhibiting GSDMD pore formation via disulfiram could provide substantial clinical benefit. In addition to improving host immune responses, disulfiram may have antiviral effects on SARS-CoV-2. Indeed, disulfiram has been shown to inhibit papain-like proteases of deadly coronaviruses such as Middle East respiratory syndrome coronavirus (MERS-CoV) and SARS-CoV-1, which may disrupt the replication and IFN suppression mechanisms of these viruses.^79–81^ Disulfiram was shown to inhibit the main protease of SARS-CoV-2 by demonstrating high inhibitory concentration values but the direct inhibition of SARS-CoV-2 viral replication by disulfiram has yet to be demonstrated.^79^ Recently, a clinical trial was initiated at the University of San Francisco to test how three days of disulfiram treatment changes adverse events, disease severity, SARS-COV-2 viral load, and inflammatory cytokine production in patients with early COVID-19 (ClinicalTrials.gov Identifier: NCT04485130).

We plan to test the most promising and safe compounds discovered by our multi-phenotype aging profile of viral infection for their ability to impact clinical care in sepsis, influenza, ARDS, and COVID-19. At the same time, we hope to further develop our system by building the capability to generate aging profiles of other age-related diseases such as autoimmune conditions, cancer, and vaccine responses. If successful, our current and future work could be used to improve the health of many older adults.

## MATERIALS AND METHODS

### Study Ethics and Donors

All study protocols were evaluated and approved by the WCG Institutional Review Board, and all donors provided informed consent. Eighty-nine healthy donors underwent screening at the Discovery Life Sciences Donor Clinic (Huntsville, AL). No donors were excluded based on comorbidities, alcohol use, smoking, or current medication.

### Sample Collection and Cell Isolation

Whole blood was collected into EDTA tubes and then diluted with an equal volume of phosphate-buffered saline (PBS) + 2% fetal bovine serum (FBS) and layered over Ficoll using SepMate™-50 tubes (STEMCELL Technologies Inc., Vancouver, Canada). Cells were centrifuged at 1200×*g* for 10 min at room temperature, and the top plasma layer was removed.

PBMCs were collected, washed with PBS + 2% FBS, and counted using acridine orange/propidium iodide using a Cellometer^®^ Vision CBA (Nexcelom Bioscience, Lawrence, MA, USA). PBMCs were cryopreserved in CryoStor^®^ CS10 (BioLife Solutions, Bothell, WA, USA), frozen using CoolCell^®^ FTS30 freezing containers (BioCision, San Rafael, CA, USA), and stored in the liquid nitrogen vapor phase until use.

Specific cell types were isolated from the PBMC fraction using the following kits (STEMCELL Technologies Inc.) per manufacturer’s recommendations: EasySep™ Human T Cell Enrichment Kit (T cells); EasySep™ Human B Cell Enrichment Kit (B cells); EasySep™ Human NK Cell Enrichment Kit (NK cells); and EasySep™ Human Monocyte Enrichment Kit (monocytes). T cells (CD3^+^), B cells (CD19^+^), NK cells (CD56^+^), and monocytes (CD14^+^) were isolated based on their customarily defined cell surface markers. Isolated cells were counted using acridine orange/propidium iodide on a Cellometer Vision CBA and then cryopreserved as described above.

### Assay Plate Layout

Each 384-well plate assay layout was designed to account for potential bias in experimental conditions due to plate location; conditions were plated such that every condition was included in every row or column. To generate data on isolated cell types, T cells, monocytes, macrophages, dendritic cells, B cells, NK cells, and PBMCs were evaluated as separate conditions. To evaluate effects due to viral load, we plated unexposed cells and cells exposed at 0.1×, 1×, and 10× MOI as different conditions (described below). To analyze differences between age groups, we plated 48 individual donors per assay plate, with 8 replicates per donor. Two plates were required to evaluate all donors. Donors were stratified by age group (≤35 vs ≥60 years). The age groups were considered as separate conditions such that the interspersed plating resulted in a balance between young and old donors across all plate locations.

### Thawing and Plating Cells

Cells were thawed from the gas phase of liquid nitrogen by immersing the cryotube in a 37 °C water bath until a thin layer of ice was present. The cells were transferred to 10 mL of prewarmed culture medium (refer to Supplemental Methods for all media recipes) and immediately centrifuged for 10 min at 138×*g*. The cell pellet was gently washed with 10 mL of culture medium, resuspended in cold plating medium at a density of 4×10^5^ cells/mL, and 25 μL were added to the assay plate (filled with 15-μL prewarmed plating medium) using a 96-well dispensing head to achieve a final cell density of 10,000 cells per well. Cells were kept cool during the entire plating process. Assay plates were centrifuged for 1 min at 138×*g* and incubated for 30 min at 37 °C with 5% CO_2_ to allow for cell adhesion to the plastic surface of the assay plate.

### Vesicular Stomatitis Virus and Compound Exposure

After 30-min cell adhesion, 10 μL of 5× trigger medium (including rVSV-ΔG-mCherry, DMSO, test compound, and FBS) was added to the assay plate using a 384-well pipetting head to achieve a final concentration of rVSV-ΔG-mCherry at different MOIs, 0.1% DMSO, 10% FBS, and 0.33 μM or 5.3 μM compound concentration. The assay plate was centrifuged for 1 min at 138×*g* and incubated for 24 h at 37 °C with 5% CO_2_.

### Live Cell Staining, Fixation, and Painting

After 24 h, cell supernatant was removed and evaluated to determine cytokine levels using the FirePlex^®^-HT assay system (Abcam, Cambridge, MA, USA; described below). To label mitochondria in live cells for Cell Painting Palette 2, 50 μL/well of prewarmed 125 nM cell-permeant MitoTracker^®^ solution (Thermo Fisher Scientific, Waltham, MA, USA) was added to the assay plate and incubated at 37 °C with 5% CO_2_. After 30 min incubation, cells were washed 2× with 50 μL/well prewarmed culture medium. Cells were fixed through replacing 25 μL of the culture medium with 25 μL of prewarmed formaldehyde solution in culture medium (8% v/v). For Cell Painting Palette 1, cells were fixed adding 50 μL of formaldehyde solution in PBS (4% v/v) directly after removal of cell supernatant. Cells were incubated at room temperature for 20 min before washing 3× with 50 μL/well PBS. Cells were then permeabilized with 50 μL/well permeabilization buffer (Supplemental Methods) for 5 min, then washed with 50 μL/well PBS. Block buffer (50 μL/well; Supplemental Methods) was added to each well, and the assay plate was incubated for 1 h at room temperature.

For the cell painting assay, Cell Painting Palette 1 or 2 (Supplemental Methods) was set up in block buffer. The palette was similar to the cell painting assay described by Bray et al.^28^ Block buffer was removed from the cells and 25 μL/well of the CPMM palette was added. After an overnight (~18 h) incubation at 8 °C, the assay plate was washed 4× with 50-μL/well PBS/ Antibiotic Antimycotic and subjected to image acquisition. The plate was stored under refrigerated conditions for further use.

### Imaging

Image acquisition was conducted on an Operetta CLS™ high-content analysis system (PerkinElmer, Waltham, MA) with binning 1, using a 40× water immersion objective in wide-field mode with Z-stack of 4 planes and distance steps of 1.0 μm. Four to 16 fields of view and six channels were captured.

### Cytokine Detection

Cytokine levels in the cellular supernatant were evaluated using the FirePlex-HT assay system with the Human Cytokines FirePlex-HT Panel 1 (ab234897; Abcam). FirePlex-HT immunoassays quantify up to 10 protein analytes per sample from low sample inputs in 384-well plate format. The following cytokines were evaluated: IFNγ, IL-1β, IL-2, IL-4, IL-6, IL-8, IL-10, IL-17A, MCP1, and TNFα.

### Flow Cytometry

Cells were plated as described above at a cell density of 1.5×10^5^ cells/cm^2^ in 24-well microtiter plates and incubated for 24 h at 37 °C and 5% CO_2_ All cells, including the floating cells, were harvested by adding 1 mL PBS/EDTA (1 mM) and incubating for 5 min at 37 °C before being transferred to a round-bottom flow cytometry tube. All remaining steps were carried out with the cells either kept on ice or at 4 °C. Cells were washed once with 5 mL of cold FACS buffer (Supplemental Methods) and centrifuged at 216×*g* for 10 min, then stained with 100-μL antibody mix for 45 min protected from light. Cells were washed 2× with 5-mL FACS buffer, resuspended in 0.5-mL PBS, passed through a cell strainer, and analyzed using a FACSAria™ I cell sorter (Becton, Dickinson and Company, Franklin Lakes, NJ, USA). For VSV-treated experiments, monocytes were defined by medium to high forward scattered light (FSC) and side scattered light (SSC) morphology and CD11c^+^ CD14^+^. NK cells were defined by low FSC/SSC CD3^−^ CD56^+^, and T cells were defined by low FSC/SSC and CD3^+^. Within each defined cellular population, infected cells were segregated by mCherry+ signal. Similar gating structures were used for initial identification of cellular subsets in the absence of VSV infection, with the addition of CD19^+^ FSC/SSC low events defined as B cells (see Supplemental Figure 1).

### Image Processing

Initial data preprocessing was structured as a directed acyclic graph, with image transformation at each step in the graph execution; transformed images were passed along as inputs to the next step. For the first step, the input images, acquired as multiple confocal Z-layers per field of view, were projected onto a single field of view by taking the maximum value of each pixel across all Z-layers for a given field. The projected images were saved both at the original source resolution and at a down-sampled half-resolution. For each channel, an illumination correction function was computed using the half-resolution images and then applied to the full-resolution and half-resolution images to generate illumination-corrected versions of the images.^82^ For each channel, the mean and standard deviation of the pixel value distributions were computed over the projected images. The full-resolution and half-resolution images were then *z*-score normalized using these precomputed means and standard deviations of the plate for each channel to generate pre-normalized versions of the images.

### Feature Generation

Using the normalized images, the data processing method continued to feature generation. A CellProfiler pipeline was generated by parameterizing on the relevant palette, magnification, and cells used for the experiment.^83^ CellProfiler uses the locations of each cell to generate over 700 features per cell, including cell morphology, stain intensity, stain texture, stain granularity, and stain colocalization across channels. The CellProfiler pipeline was run over the full-resolution, illumination-corrected images to segment and generate thousands of image- and cell-level features. The location of each cell and nucleus on the plate, as well as a TIFF-based mask for each, was extracted from the CellProfiler output and converted to a format amenable to downstream processing. Pretrained convolutional neural networks were then used to extract single-cell deep-learning embeddings for every cell detected by CellProfiler in every channel. Embeddings were generated at multiple output layers in the pretrained model and then concatenated to create a single embedding vector for each cell that includes 3840 features. The individual cells were fed to the model using 88px single cell crops taken from the prenormalized 2160×2160 pixel images. For specific steps in the CellProfiler pipeline, please refer to the Supplemental Methods.

### Quality Control

Every image, and each cell within the image, was run through a semi-automated quality control process. The image metrics computed by CellProfiler were visualized and analyzed per plate to identify technical artifacts that arose from issues such as location, operator error, and poor sample quality (Supplemental Figure S4). Each plate was reviewed by an operator to check for quality across cell count, intensity, blur, focus, and outlier metrics. Plates that had strong negative or positive correlations between quality control metrics and plate location were marked for further review. These metrics were used to flag problematic issues, refine assay protocols, and inform interpretation of results, but at this point were not used to exclude data in an automatic fashion.

### Spatial Point Pattern Analysis

The individual cell points detected by CellProfiler were analyzed using spatial point pattern analysis. This modeling provides a statistical basis for understanding the effect of different conditions on how cells spatially interact (clustering or inhibitory) with other cells. After processing, individual fields were treated as unique spatial windows. The point locations of individual cells were marked with labels within these windows, effectively translating an image of a field to a marked point pattern. A data set was then constructed using these marked point patterns, relevant spatial covariates (eg, relative field location), and global covariates (eg, donor identification and well condition). This marked point pattern data set was used to model the underlying point pattern process. Spatial point patterns of individual fields were modeled with non-Poisson multitype point interactions for specific cell types. From these individual point pattern models, cell-to-cell interaction parameters for specific label-to-label interactions (eg, T cell to macrophage) were calculated for each field for subsequent analysis. Spatial point patterns of a selection of fields were modeled using replicated spatial point pattern modeling. An additional series of stepwise replicated point pattern models with non-Poisson multitype point interactions were constructed to analyze point pattern interactions of different cell types with respect to conditions of interest. The replicated point pattern models could include a number of global and spatial covariates to model heterogeneous and homogeneous point patterns, varying interaction parameters describing interpoint interactions, and interactive interaction parameters describing the relationship of conditions of interest and cell-to-cell interactions. Interaction scores <1 indicated decreased clustering while interaction scores >1 indicated increased clustering. This modeling provides evidence of whether or not the conditions of interest alter the cell-to-cell interactions of targeted labels (eg, monocyte to monocyte and T cell to macrophage), as well as how cell-to-cell interactions change with these conditions (eg, T cells exhibiting increased clustering with macrophages under a given condition).

### Classification

We used the embeddings and features created to train classifiers that predicted whether an immune response was indicative of a young or old immune system. Data-generation methods were optimized to reduce bias by using balanced plate layouts, robust quality control processes, automated liquid handling, and a large, diverse number of samples. Embeddings were generated for each single cell and then aggregated by taking the median of all cells within a field of view. We trained the classification models on these field-level embeddings using XGBoost, an implementation of gradient-boosted decision trees.^72^ As described in the Results, we performed stratified four-fold cross-validation, ensuring that each fold had equal proportions of young and old donors, and that each donor appeared in only one-fold.

## ACKNOWLEDGEMENTS

Acquisition of data was provided by Assay.Works GmbH (Regensburg, Germany), funded by Spring Discovery. Sample collection and processing was provided by Discovery Life Sciences (Huntsville, AL, USA), funded by Spring Discovery. Medical writing and/or editorial assistance was provided by Rebecca Brady, PhD, of The Lockwood Group (Stamford, CT, USA), funded by Spring Discovery Inc.

## AUTHOR CONTRIBUTIONS

**Experimental conception and design**: Brandon White, Ben Komalo, Lauren Nicolaisen, Matt Donne, Charlie Marsh, Rachel M. DeVay, An M. Nguyen, Wendy Cousin, Jarred Heinrich, William Van Trump, Colin J. Fuller, Laura Haynes, George Kuchel, Jorg Goronzy, Tamas Fulop, Diane Heiser, Ralf Schwandner, Christian Elabd, Ben Kamens. **Data acquisition**: Brandon White, Matt Donne, William Van Trump, Tempest Plott, Dat Nguyen, Daniel Chen, Delia Bucher, Sabine Tyrra, Ralf Schwandner. **Data analysis**: Brandon White, Ben Komalo, Lauren Nicolaisen, Charlie Marsh, Rachel M. DeVay, An M. Nguyen, Wendy Cousin, Jarred Heinrich, Diane Heiser. **Data interpretation**: Brandon White, Ben Komalo, Lauren Nicolaisen, Charlie Marsh, Rachel M. DeVay, An M. Nguyen, Wendy Cousin, Jarred Heinrich, Colin J. Fuller, Laura Haynes, George Kuchel, Jorg Goronzy, Anis Larbi, Tamas Fulop, Diane Heiser, Ralf Schwandner, Christian Elabd. **Software creation**: Ben Komalo, Lauren Nicolaisen, Charlie Marsh, Jarred Heinrich, Colin J. Fuller. **Supervision of research**: Brandon White, Ralf Schwandner, Christian Elabd, Ben Kamens. **Drafting and editing the manuscript**: Brandon White. All authors revised and approved the final manuscript.

## COMPETING INTERESTS

Brandon White, Ben Komalo, Lauren Nicolaisen, Matt Donne, Charlie Marsh, Rachel M. DeVay, An M. Nguyen, Wendy Cousin, Jarred Heinrich, William Van Trump, Tempest Plott, Colin J. Fuller, Dat Nguyen, Daniel Chen, Christian Elabd, and Ben Kamens are employees of, and have an equity interest in, Spring Discovery. Brandon White, Ben Komalo, Lauren Nicolaisen, Charlie Marsh, An M. Nguyen, Wendy Cousin, Christian Elabd, and Ben Kamens are co-inventors on patents filed for this work and pending to Spring Discovery. Laura Haynes and Jorg Goronzy report grants from NIH. Laura Haynes, George Kuchel, Jorg Goronzy, Anis Larbi, and Ralf Schwandner report personal fees from Spring Discovery. Anis Larbi is the co-founder of Yobitrust. Ralf Schwandner is the founder and CEO of Assay.Works GmBH. Delia Bucher, Sabine Tyrra, Tamas Fulop, and Diane Heiser have nothing to disclose.

## SUPPLEMENTAL INFORMATION

## SUPPLEMENTAL FIGURES

**Supplemental Figure S1.**
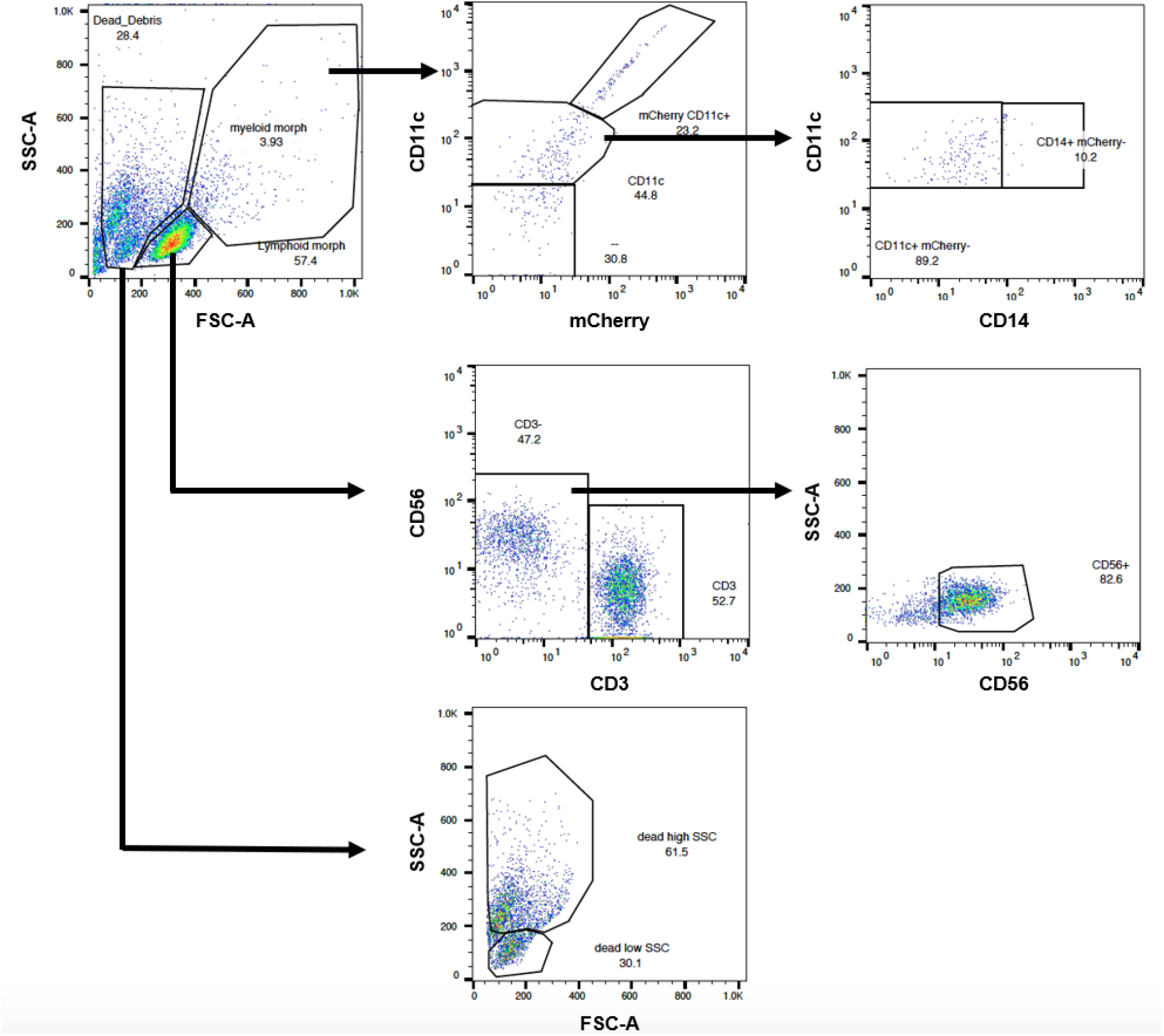
Representative flow plots for cell composition discrimination. Events were first gated on FSC/SSC profile to distinguish dead cells and debris from live myeloid and live lymphoid cells. Myeloid morphology cells were then stratified by CD14 and CD11c expression to distinguish monocytes (CD14^+^). Lymphoid cells were first stratified by CD3 and CD19 expression to distinguish T cells (CD3^+^) and B cells (CD19^+^). Double-negative cells from the lymphoid morphology fraction were used to distinguish NK cells (CD3^−^ CD19^−^ CD56^+^).

**Supplemental Figure S2.**
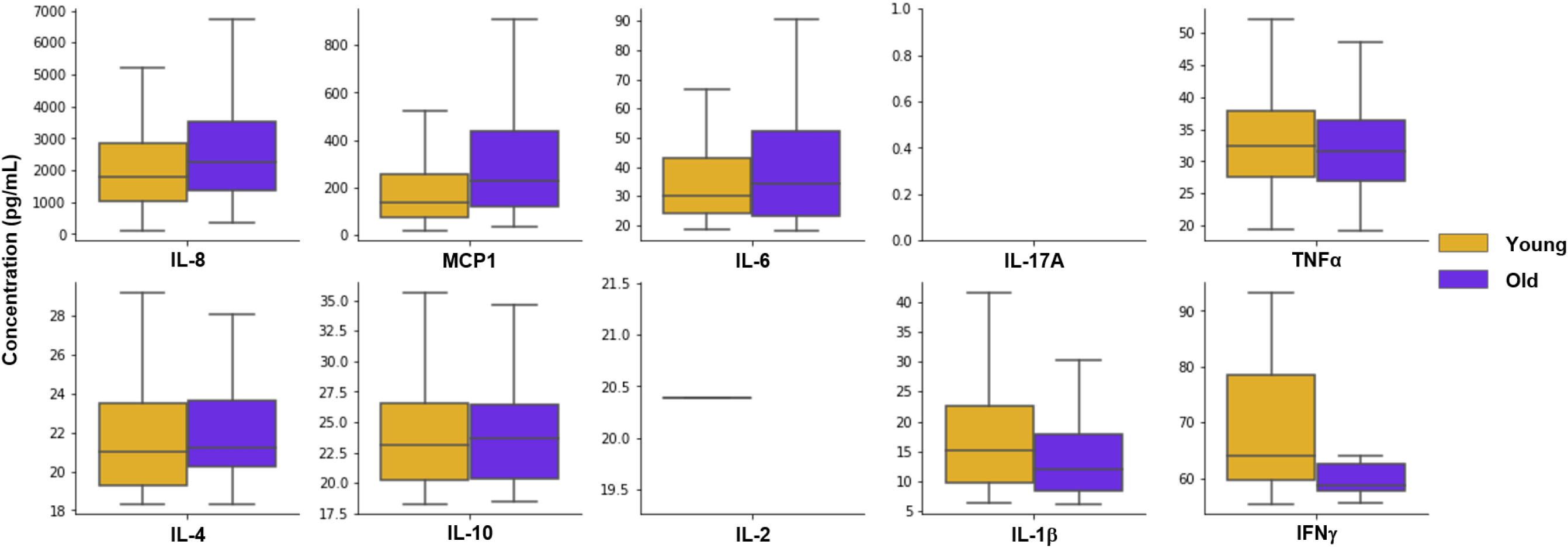
Cytokine levels from the supernatant of PBMCs obtained from 89 old and young donors and exposed to rVSV-ΔG-mCherry at 1× MOI. IFN, interferon; IL, interleukin; MCP, monocyte chemoattractant protein 1; MOI, multiplicity of infection; PBMC, peripheral blood mononuclear cell; rVSV-ΔG-mCherry, recombinant vesicular stomatitis virus expressing a red fluorescent construct; TNF, tumor necrosis factor.

**Supplemental Figure S3.**
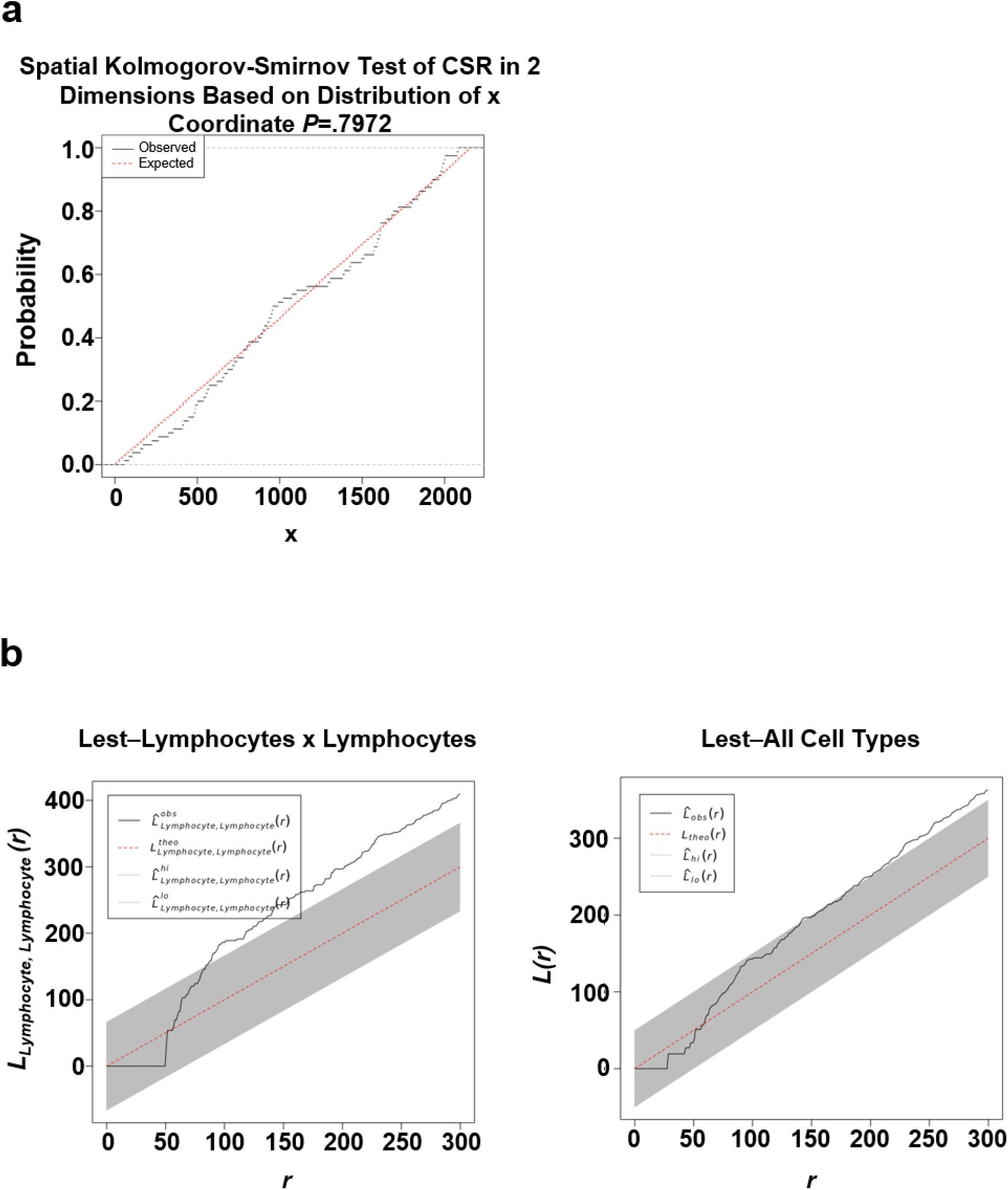
(a) Spatial Kolmogorov-Smirnov plot to identify inconsistencies with a homogeneous Poisson distribution. (b) Significant departures from the expected distribution (homogeneous) would indicate a heterogeneous distribution that is not independent of the spatial landscape. CSR, complete spatial randomness.

**Supplemental Figure S4.**
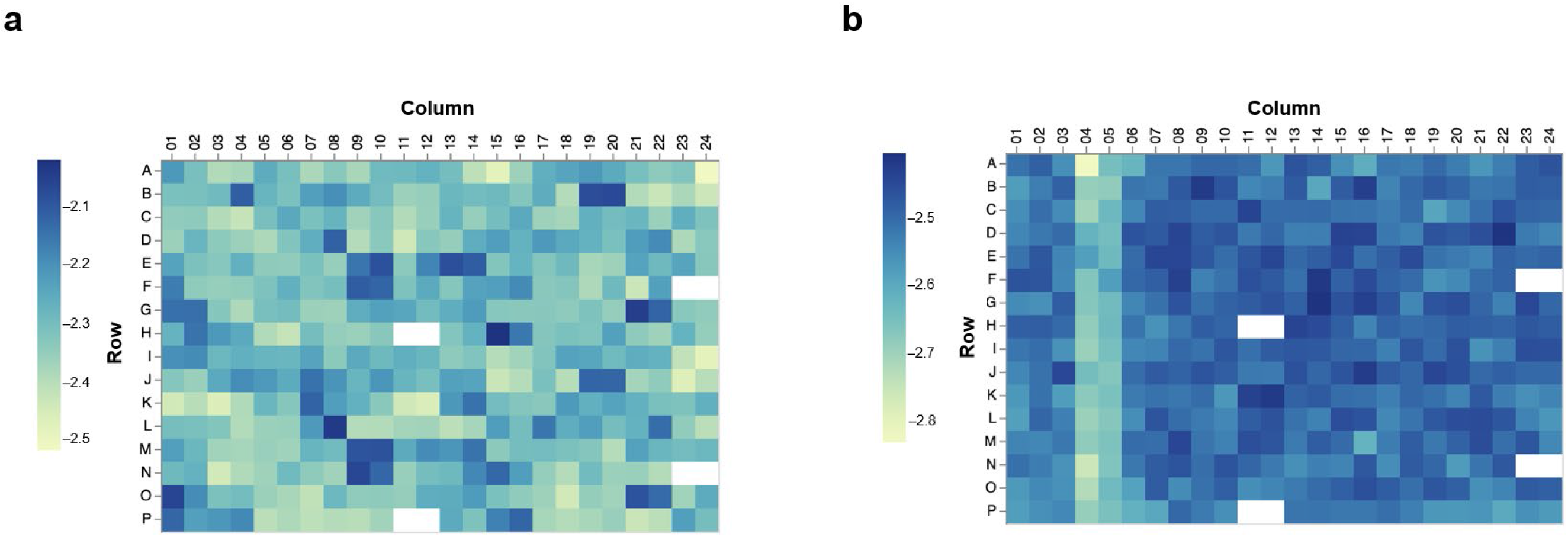
Power log-log metrics computed on the Hoechst channel for (a) a plate that passed quality control and (b) a plate flagged for quality control due to technical artifacts.

**Supplemental Figure S5.**
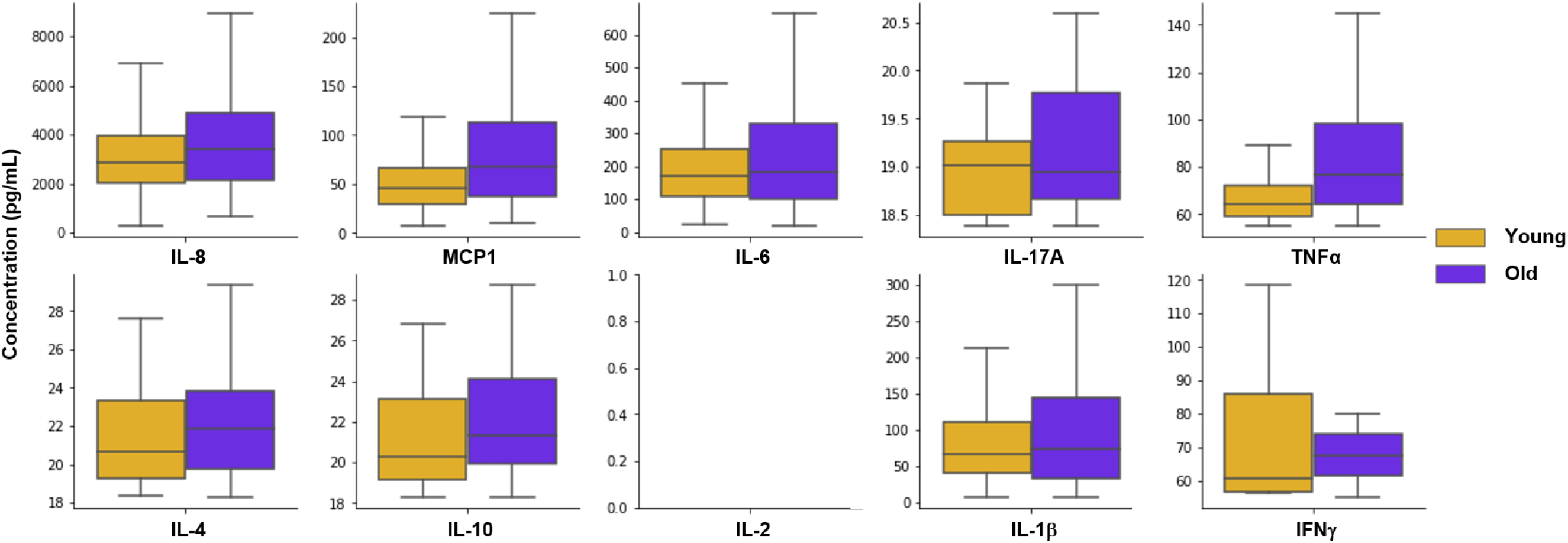
Replicate experiment measuring cytokine levels from the supernatant of rVSV-ΔG-mCherry-exposed (at 10× MOI for 24 h) PBMCs obtained from 89 donors from young and old adult populations.

**Supplemental Table S1.** Cell Painting Palette 1

| Channel | Label | Excitation (nm) | Emission (nm) | Stain |
| --- | --- | --- | --- | --- |
| 1 | Brightfield | Transmission | 655–760 |  |
| 2 | Hoechst 33342 | 355–385 | 430–500 | Hoechst |
| 3 | Alexa 488 | 460–490 | 500–550 | Wheat germ agglutinin |
| 4 | Texas Red | 530–560 | 570–650 | Phalloidin |
| 5 | Alexa 633 | 615–645 | 655–705 | Concanavalin A |

**Supplemental Table S2.** Cell Painting Palette 2

| Channel | Label | Excitation (nm) | Emission (nm) | Stain |
| --- | --- | --- | --- | --- |
| 1 | Brightfield | Transmission | 655–760 |  |
| 2 | Hoechst 33342 | 355–385 | 430–500 | Hoechst |
| 3 | Alexa 488 | 460–490 | 500–550 | Wheat germ agglutinin |
| 4 | MitoTracker Orange | 530–560 | 570–650 | MitoTracker/mCherry |
| 5 | Alexa 633 | 615–645 | 655–705 | Concanavalin A |
| 6 | Alexa 700_narrow | 650–675 | 685–760 | Phalloidin iFluor700 |

**Supplemental Table S3.**
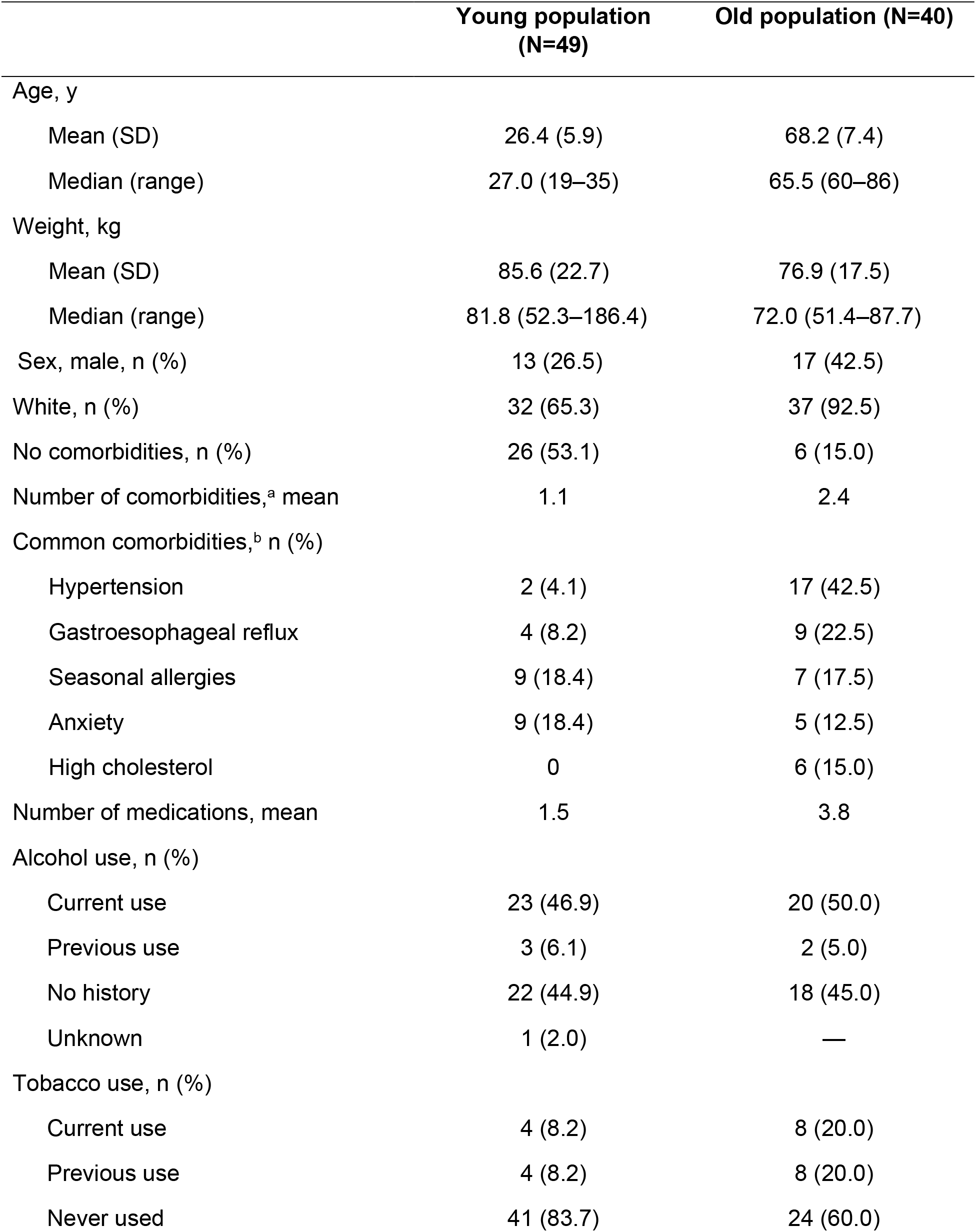
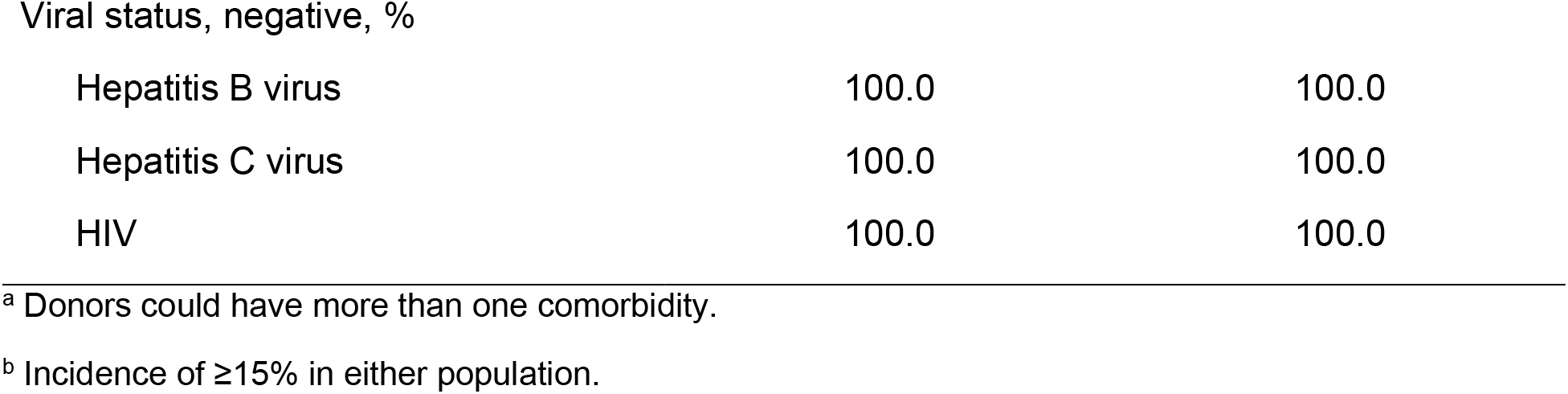
Demographics of Sample Donors

|  | Young population<br>(N=49) | Old population (N=40) |
| --- | --- | --- |
| Age, y |  |  |
| Mean (SD) | 26.4 (5.9) | 68.2 (7.4) |
| Median (range) | 27.0 (19–35) | 65.5 (60–86) |
| Weight, kg |  |  |
| Mean (SD) | 85.6 (22.7) | 76.9 (17.5) |
| Median (range) | 81.8 (52.3–186.4) | 72.0 (51.4–87.7) |
| Sex, male, n (%) | 13 (26.5) | 17 (42.5) |
| White, n (%) | 32 (65.3) | 37 (92.5) |
| No comorbidities, n (%) | 26 (53.1) | 6 (15.0) |
| Number of comorbidities,<sup>a</sup> mean | 1.1 | 2.4 |
| Common comorbidities,<sup>b</sup> n (%) |  |  |
| Hypertension | 2 (4.1) | 17 (42.5) |
| Gastroesophageal reflux | 4 (8.2) | 9 (22.5) |
| Seasonal allergies | 9 (18.4) | 7 (17.5) |
| Anxiety | 9 (18.4) | 5 (12.5) |
| High cholesterol | 0 | 6 (15.0) |
| Number of medications, mean | 1.5 | 3.8 |
| Alcohol use, n (%) |  |  |
| Current use | 23 (46.9) | 20 (50.0) |
| Previous use | 3 (6.1) | 2 (5.0) |
| No history | 22 (44.9) | 18 (45.0) |
| Unknown | 1 (2.0) | — |
| Tobacco use, n (%) |  |  |
| Current use | 4 (8.2) | 8 (20.0) |
| Previous use | 4 (8.2) | 8 (20.0) |
| Never used | 41 (83.7) | 24 (60.0) |
| Viral status, negative, % |  |  |
| Hepatitis B virus | 100.0 | 100.0 |
| Hepatitis C virus | 100.0 | 100.0 |
| HIV | 100.0 | 100.0 |

<sup>a</sup> Donors could have more than one comorbidity.
<sup>b</sup> Incidence of $\geq 15\%$ in either population.

## SUPPLEMENTAL METHODS

### CellProfiler Pipeline

1. Segmenting the nuclei using the Hoechst channel, via Otsu or minimum cross entropy-based thresholding
2. Segmenting the cell membrane using an appropriate channel, via minimum cross entropy-based thresholding
3. Computing, over both the nuclei and cell membranes, a set of features to capture:

a. The size and shape of the objects of interest
b. The texture of the objects of interest, across all channels
c. The stain colocalization of the objects of interest, across all pairs of channels
d. The granularity of the objects of interest, across all channels
e. The intensity of the staining of the objects of interest, across all channels
f. The intensity distribution of the staining of the objects of interest, across all channels
g. A TIFF-based mask of the object
4. Computing, over the entire field of view:

a. The intensity of the staining, across all channels
b. The image correlation, power log-log slope, and other quality control metrics, across all channels

### Materials and Equipment

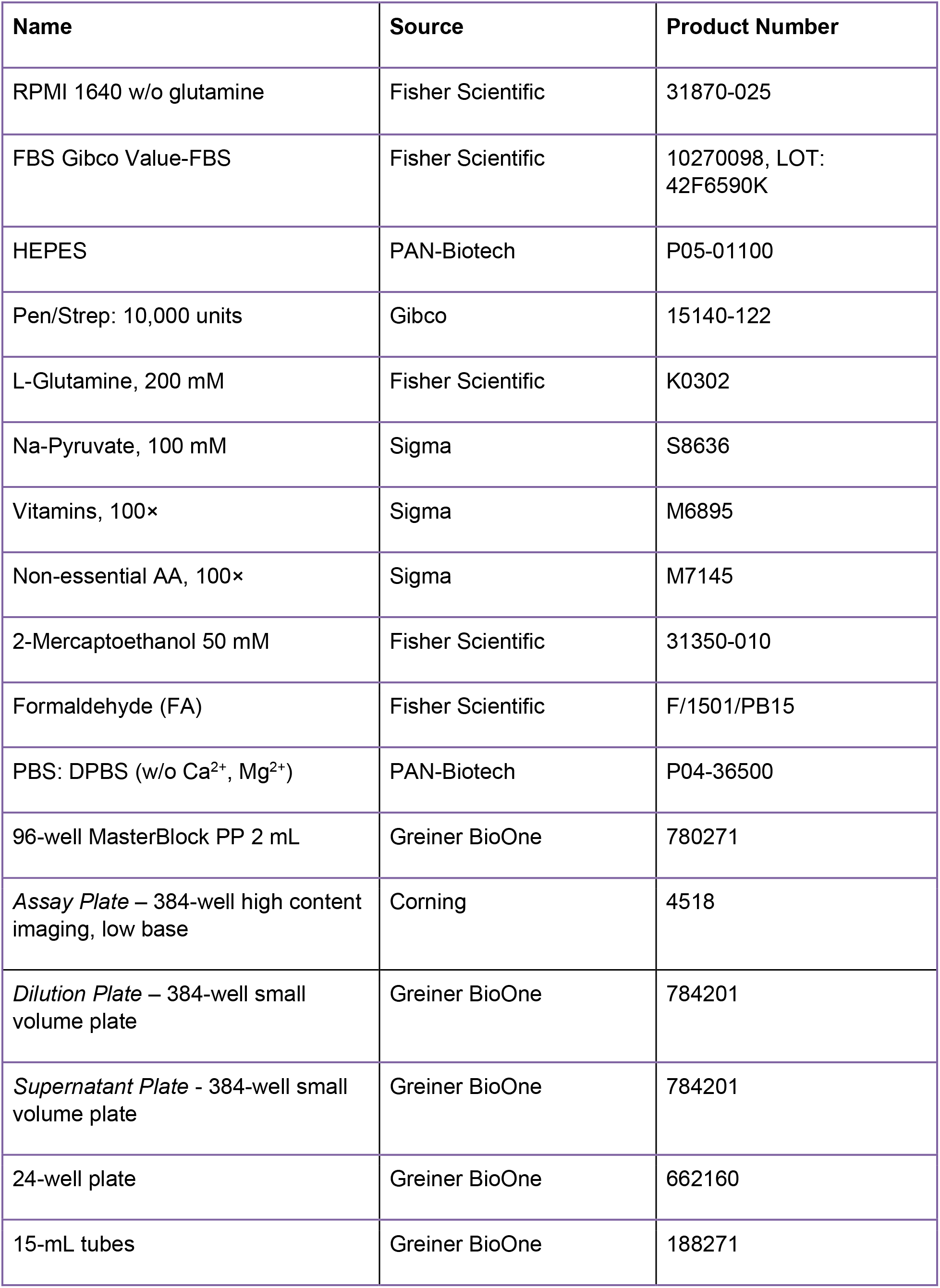
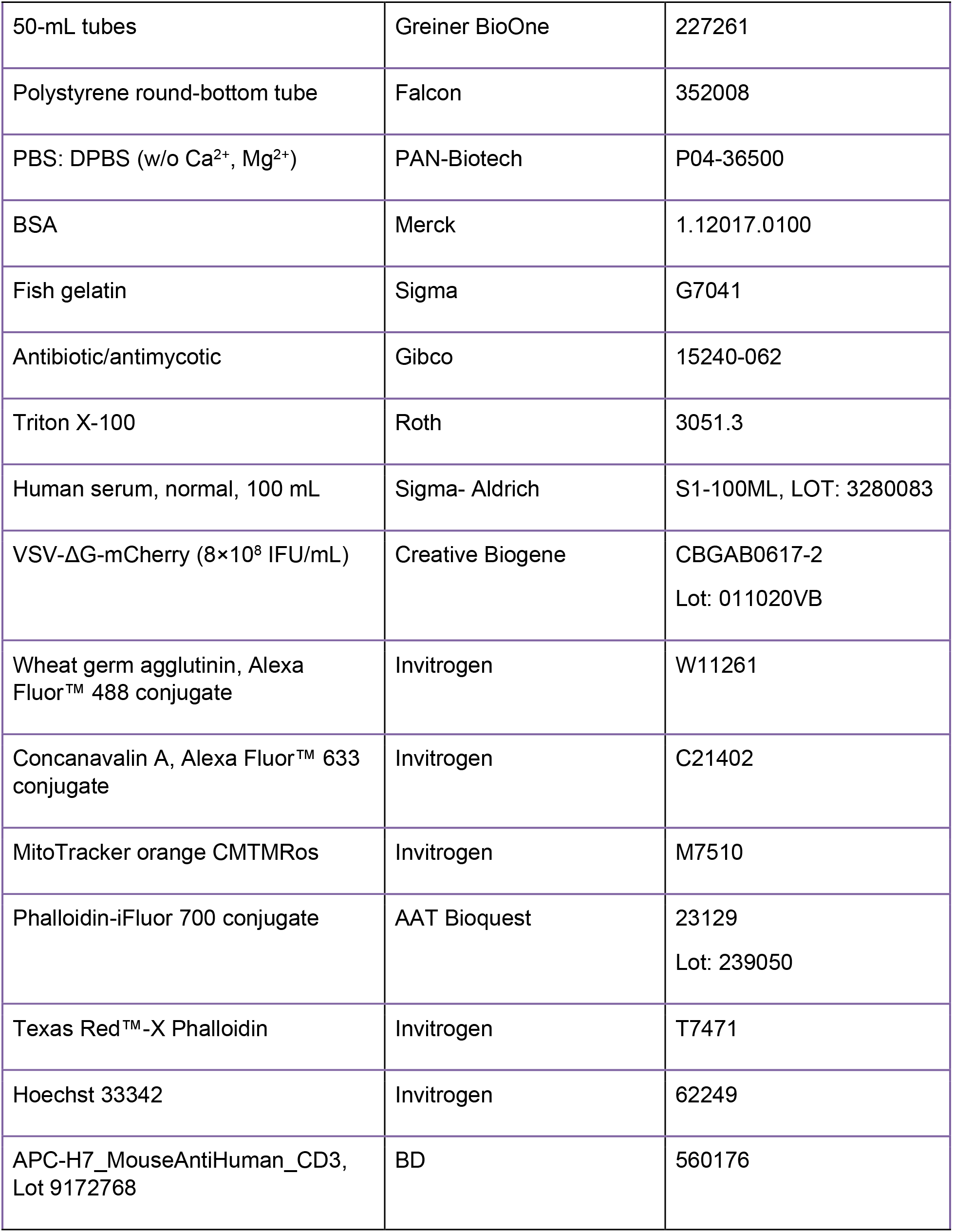
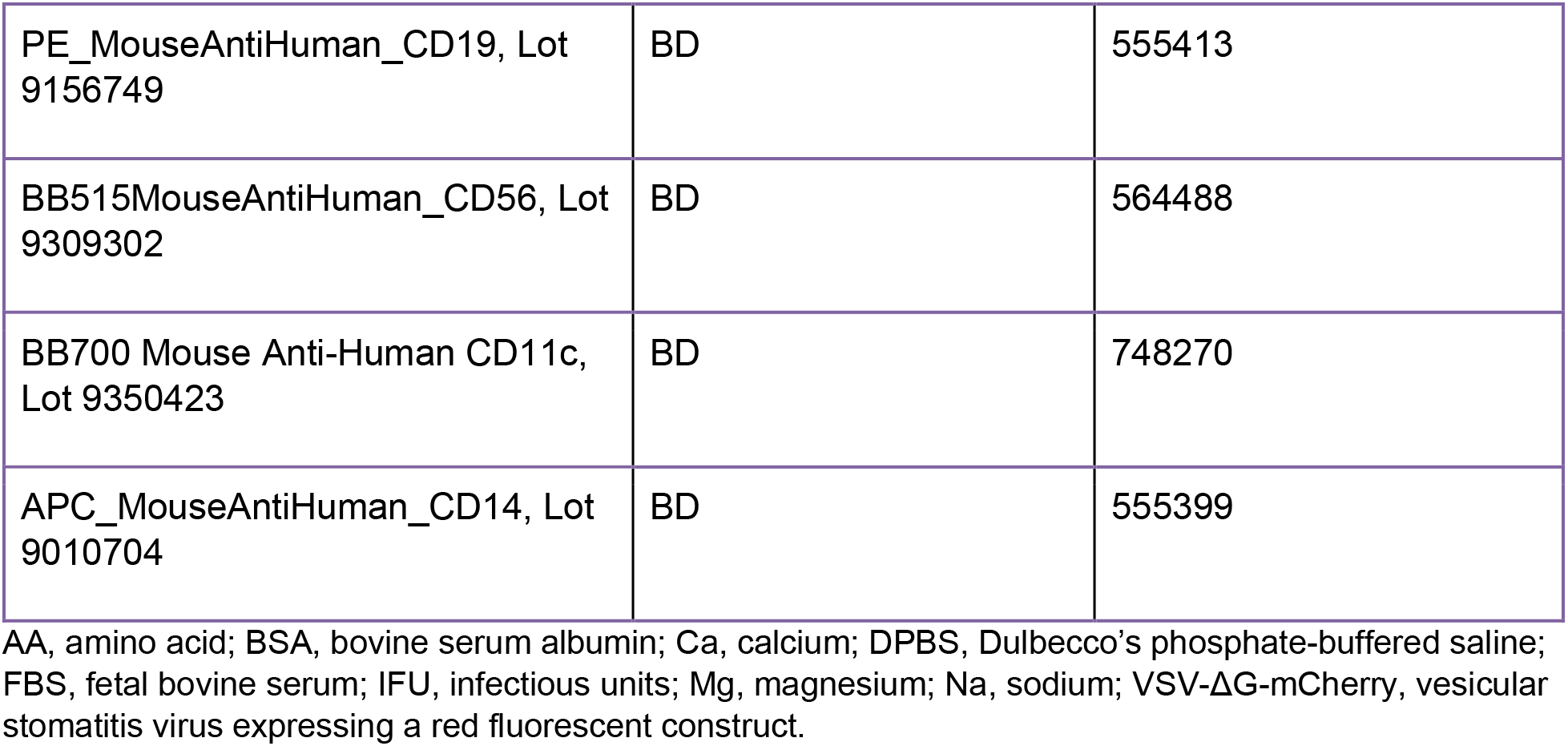

| **Name** | **Source** | **Product Number** |
| --- | --- | --- |
| RPMI 1640 w/o glutamine | Fisher Scientific | 31870-025 |
| FBS Gibco Value-FBS | Fisher Scientific | 10270098, LOT:<br>42F6590K |
| HEPES | PAN-Biotech | P05-01100 |
| Pen/Strep: 10,000 units | Gibco | 15140-122 |
| L-Glutamine, 200 mM | Fisher Scientific | K0302 |
| Na-Pyruvate, 100 mM | Sigma | S8636 |
| Vitamins, 100× | Sigma | M6895 |
| Non-essential AA, 100× | Sigma | M7145 |
| 2-Mercaptoethanol 50 mM | Fisher Scientific | 31350-010 |
| Formaldehyde (FA) | Fisher Scientific | F/1501/PB15 |
| PBS: DPBS (w/o Ca<sup>2+</sup>, Mg<sup>2+</sup>) | PAN-Biotech | P04-36500 |
| 96-well MasterBlock PP 2 mL | Greiner BioOne | 780271 |
| *Assay Plate* – 384-well high content imaging, low base | Corning | 4518 |
| *Dilution Plate* – 384-well small volume plate | Greiner BioOne | 784201 |
| *Supernatant Plate* - 384-well small volume plate | Greiner BioOne | 784201 |
| 24-well plate | Greiner BioOne | 662160 |
| 15-mL tubes | Greiner BioOne | 188271 |
| 50-mL tubes | Greiner BioOne | 227261 |
| Polystyrene round-bottom tube | Falcon | 352008 |
| PBS: DPBS (w/o Ca<sup>2+</sup>, Mg<sup>2+</sup>) | PAN-Biotech | P04-36500 |
| BSA | Merck | 1.12017.0100 |
| Fish gelatin | Sigma | G7041 |
| Antibiotic/antimycotic | Gibco | 15240-062 |
| Triton X-100 | Roth | 3051.3 |
| Human serum, normal, 100 mL | Sigma- Aldrich | S1-100ML, LOT: 3280083 |
| VSV-ΔG-mCherry (8×10<sup>8</sup> IFU/mL) | Creative Biogene | CBGAB0617-2<br>Lot: 011020VB |
| Wheat germ agglutinin, Alexa Fluor™ 488 conjugate | Invitrogen | W11261 |
| Concanavalin A, Alexa Fluor™ 633 conjugate | Invitrogen | C21402 |
| MitoTracker orange CMTMRos | Invitrogen | M7510 |
| Phalloidin-iFluor 700 conjugate | AAT Bioquest | 23129<br>Lot: 239050 |
| Texas Red™-X Phalloidin | Invitrogen | T7471 |
| Hoechst 33342 | Invitrogen | 62249 |
| APC-H7_MouseAntiHuman_CD3,<br>Lot 9172768 | BD | 560176 |
| PE_MouseAntiHuman_CD19, Lot 9156749 | BD | 555413 |
| BB515MouseAntiHuman_CD56, Lot 9309302 | BD | 564488 |
| BB700 Mouse Anti-Human CD11c, Lot 9350423 | BD | 748270 |
| APC_MouseAntiHuman_CD14, Lot 9010704 | BD | 555399 |

AA, amino acid; BSA, bovine serum albumin; Ca, calcium; DPBS, Dulbecco's phosphate-buffered saline; FBS, fetal bovine serum; IFU, infectious units; Mg, magnesium; Na, sodium; VSV-ΔG-mCherry, vesicular stomatitis virus expressing a red fluorescent construct.

### Specialized Materials and Equipment

| Type | Name, vendor |
| --- | --- |
| Felix - 384-well liquid handler | Cybi-Felix w/ 384-well head, Analytik Jena |
| Echo550 | Labcyte |
| Operetta CLS | PerkinElmer |
| FACS | FACS Arial Sorter, Becton Dickinson |
| Multichannel dispenser | Multidrop Combi, Thermo Fisher Scientific |
| Plate shaker | Oribital shaker GLF, digital 3017 |
| Centrifuge | Thermo Multifuge 3L-R |
| Bioactive library | Assay.Works |

FACS, fluorescence-activated cell sorting.

### Recipes

| Stock solution | Preparation | Storage; usage |
| --- | --- | --- |
| MitoTracker orange CMTMRos | Dissolved in DMSO to 1 mM | Stored at –20 °C |
| Wheat germ agglutinin, Alexa Fluor™ 488 conjugate 5 mg | Dissolved in PBS to 1 mg/mL | Single use stored at –20 °C |
| Concanavalin A, Alexa Fluor™ 633 conjugate | Dissolved in 0.1-M sodium bicarbonate (pH 8.3) to 1 mg/mL | Single use stored at –20 °C |
| Phalloidin-iFluor 700 conjugate Lot 239050 | Dissolved in 30-µL DMSO to 1000× | Stored at –20 °C |
| Texas Red™-X Phalloidin | Dissolved in 1.5 mL methanol to 200 units/mL | Stored at –20 °C |
| rVSV-ΔG-mCherry | 8×10<sup>8</sup> IFU/mL | Stored at –80 °C |
| Bioactive library | 10 mM in DMSO |  |

IFU, infectious units; PBS, phosphate-buffered saline; rVSV-ΔG-mCherry, recombinant vesicular stomatitis virus expressing a red fluorescent construct.

| Culture medium |  |
| --- | --- |
| Component | Concentration |
| RPMI 1640 | Base medium |
| FBS | 10% |
| HEPES | 25 mM |
| L-Glutamine, 200 mM | 2 mM |
| Na-Pyruvate, 100 mM | 1 mM |
| Vitamins, 100× | 0.4× |
| Non-essential AA, 100× | 1× |
| 2-Mercaptoethanol 50 mM | 50 μM |
| Pen/Strep: 10,000 units | 0.5× |

AA, amino acid; FBS, fetal bovine serum; Na, sodium.

| **Plating medium** |  |
| --- | --- |
| **Component** | **Concentration** |
| RPMI 1640 | Base medium |
| HEPES | 25 mM |
| L-Glutamine, 200 mM | 2 mM |
| Na-Pyruvate, 100 mM | 1 mM |
| Vitamins, 100× | 0.4× |
| Non-essential AA, 100× | 1× |
| 2-Mercaptoethanol 50 mM | 50 μM |
| Pen/Strep: 10,000 units | 0.5x |

AA, amino acid; FBS, fetal bovine serum; Na, sodium.

|  |
| --- |
| **5× Trigger medium<sup>a</sup>** |

| Component | Concentration |
| --- | --- |
| FBS Gibco Value-FBS | 50% |
| rVSV-ΔG-mCherry | 50× MOI ($1 \times 10^7$ IFU/mL) |
| DMSO | 0.5% (accounting for compound DMSO) |

FBS, fetal bovine serum; IFU, infectious units; MOI, multiplicity of infection; rVSV-ΔG-mCherry, recombinant vesicular stomatitis virus expressing a red fluorescent construct.
<sup>a</sup> Medium is filtered using 0.2-μm filter cups to eliminate precipitates present in the FBS.

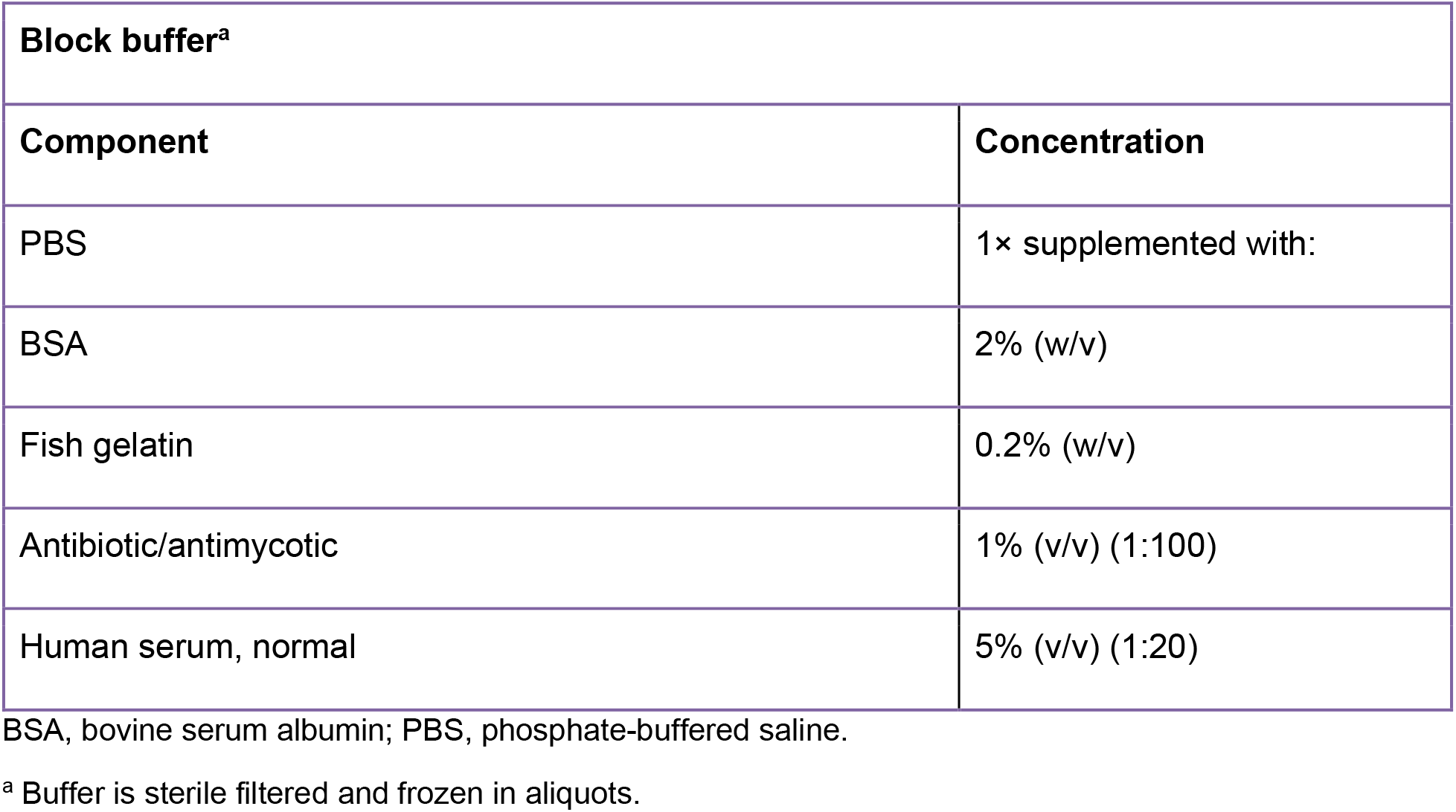

| Wash buffer |  |
| --- | --- |
| Component | Concentration |
| PBS | 1× supplemented with: |
| Antibiotic/antimycotic | 1% (v/v) (1:100) |

PBS, phosphate-buffered saline.

| **Cell Painting Palette 1<sup>a</sup>** |  |
| --- | --- |
| **Component** | **Dilution** |
| Wheat germ agglutinin, Alexa Fluor™ 488 conjugate | 1:2,000 |
| Texas Red™-X Phalloidin | 1:400 |
| Concanavalin A, Alexa Fluor™ 633 conjugate | 1:200 |
| Hoechst 33342 | 1:10,000 |

<sup>a</sup> Diluted in block buffer.

| **Cell Painting Palette 2<sup>a</sup>** |  |
| --- | --- |
| **Component** | **Dilution** |
| Wheat germ agglutinin, Alexa Fluor™ 488 conjugate | 1:2,000 |
| Phalloidin iFluor 700 prediluted stock (1:10 in DMSO) | 1:10,000 |
| Concanavalin A, Alexa Fluor™ 633 conjugate | 1:200 |
| Hoechst 33342 | 1:10,000 |

<sup>a</sup> Diluted in block buffer.

| **FACS buffer** |  |
| --- | --- |
| **Component** | **Concentration** |
| DPBS | Supplemented with |
| Human serum | 5% |

DPBS, Dulbecco's phosphate-buffered saline; FACS, fluorescence-activated cell sorting.

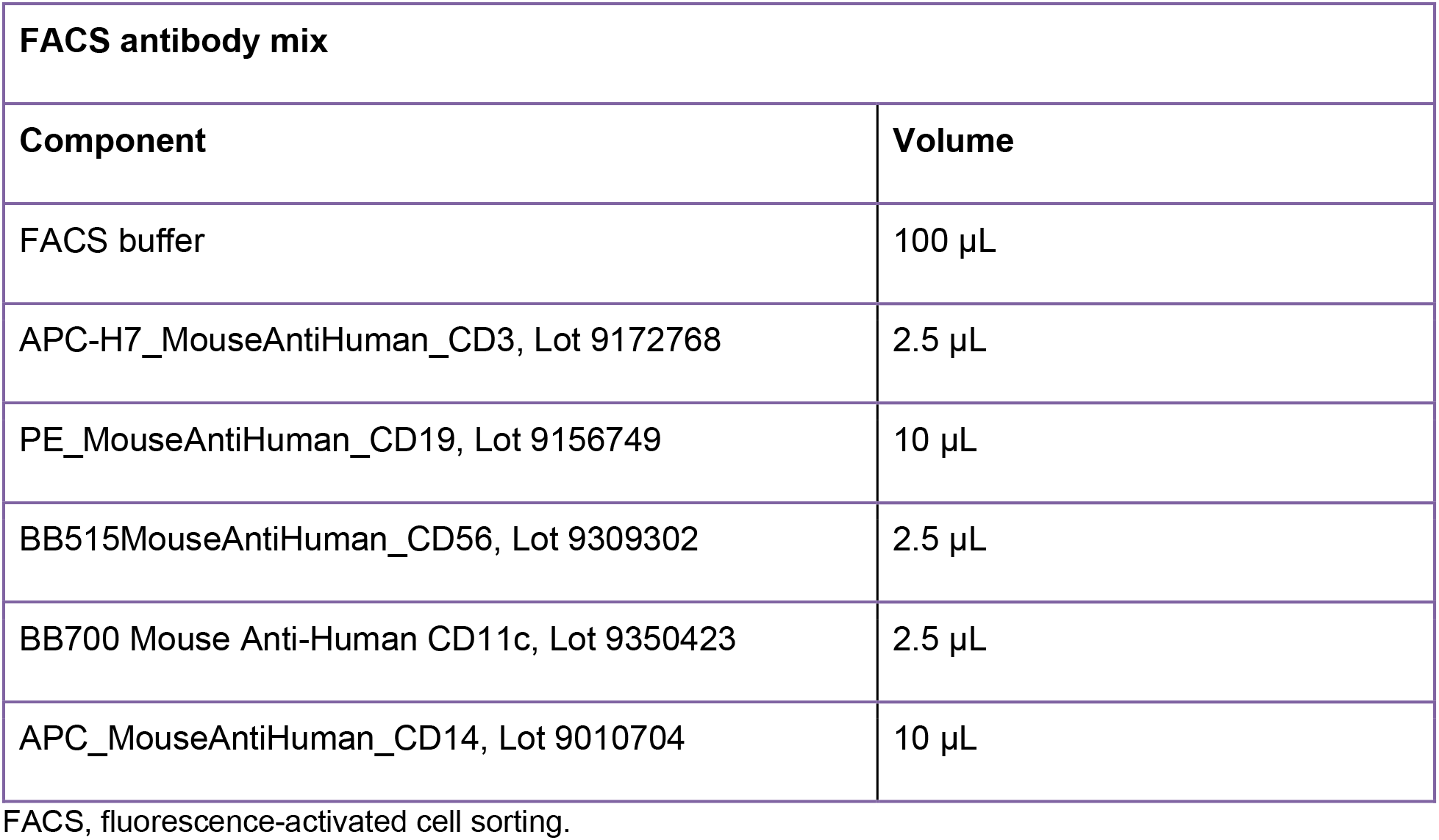

### General Experimental Notes

- Fetal bovine serum (FBS) was heat inactivated at 56 °C for 30 min before use and stored frozen in small aliquots. An aliquot was thawed for each prepared bottle of medium
- Medium was prepared fresh on the day of plating
- Trigger medium was filtered to eliminate precipitations in FBS
- Unless indicated, 5-mL serological pipettes were used throughout the protocol to minimize cell death
- 5× Trigger medium contained 5× DMSO, 5× rVSV-ΔG-mCherry, 5× test compound, and 5× FBS to obtain a final DMSO concentration of 0.1%, rVSV-ΔG-mCherry at 10× MOI, 0.33 μM or 5.3 μM compound, and 10% FBS
- All centrifugation steps were performed at room temperature (22 °C)
- Thawing and wash steps were performed with warm culture medium (37 °C)

